# Quantitative susceptibility mapping of carotid arterial tissue *ex vivo*: assessing sensitivity to vessel microstructural composition

**DOI:** 10.1101/2021.02.15.431261

**Authors:** Alan J. Stone, Brooke Tornifoglio, Robert D. Johnston, Karin Shmueli, Christian Kerskens, Caitríona Lally

**Affiliations:** Trinity Centre for Biomedical Engineering, Trinity Biomedical Sciences Institute, Trinity College Dublin, Dublin, Ireland; Department of Mechanical, Manufacturing and Biomedical Engineering, School of Engineering, Trinity College Dublin, Dublin, Ireland; Department of Medical Physics and Biomedical Engineering, University College London, London, United Kingdom; Centre for Medical Imaging, University College London, London, United Kingdom; Trinity College Institute of Neuroscience, Trinity College Dublin, Dublin, Ireland; Advanced Materials and Bioengineering Research Centre (AMBER), Royal College of Surgeons in Ireland and Trinity College Dublin, Dublin, Ireland

## Abstract

**Purpose:** To characterise microstructural contributions to the magnetic susceptibility of carotid arteries.

**Method:** Arterial vessels were scanned using high resolution quantitative susceptibility mapping (QSM) at 7T. Models of vessel degradation were generated using *ex vivo* porcine carotid arteries that were subjected to several different enzymatic digestion treatments that selectively removed microstructural components (smooth muscle cells, collagen and elastin). Magnetic susceptibilities measured in these tissue models were compared to those in untreated (native) porcine arteries. Magnetic susceptibility measured in native porcine carotid arteries was further compared to the susceptibility of cadaveric human carotid arteries to investigate their similarity.

**Results:** The magnetic susceptibility of native porcine vessels was diamagnetic (𝒳_native_ = −0.1820ppm), with higher susceptibilities in all models of vessel degradation (𝒳_elastin degraded_ = −0.0163ppm; 𝒳_collagen degraded_ = −0.1158ppm; 𝒳_decellularised_ = −0.1379ppm; 𝒳_fixed native_ = −0.2199ppm). Magnetic susceptibility was significantly higher in *collagen degraded* compared to *native* porcine vessels (Tukey-Kramer, p<0.01) and between *elastin degraded* and all other models (including *native*, Tukey-Kramer, p<0.001). The susceptibility of fixed healthy human arterial tissue was diamagnetic and no significant difference was found between fixed human and fixed porcine arterial tissue susceptibilities (ANOVA, p>0.05).

**Conclusions:** Magnetic susceptibility measured using QSM is sensitive to the microstructural composition of arterial vessels – most notably to collagen. The similarity of human and porcine arterial tissue susceptibility values provides a solid basis for translational studies. As vessel microstructure becomes disrupted during the onset and progression of carotid atherosclerosis, QSM has the potential to provide a sensitive and specific marker of vessel disease.

## 1. Introduction

### 1.1 Carotid atherosclerosis and stroke

Atherosclerosis is a widespread form of cardiovascular disease that causes the formation of plaque inside arterial vessels. Plaque formation is caused by a build-up of fat, cholesterol, calcium, fibrous tissue and other substances that disrupt the microstructural composition of the vessel, causing narrowing of the luminal space and stiffening of the vessel. Carotid arteries are particularly prone to plaque formation, and stenosis or rupture here can have catastrophic consequences as these vessels carry the main supply of blood to the brain. An estimated 20% of ischaemic stroke is due to rupture of carotid plaques, with the majority of ischaemic stroke caused by stenosis^1,2^. Carotid artery disease is also responsible for nearly 50% of transient ischaemic attacks (TIA) and linked to an increased risk of heart attack^3^. Early identification and treatment is of utmost importance to avoid stroke related disability and death caused by carotid plaque. The current indicator for surgical intervention is to assess the degree of stenosis caused by atherosclerotic plaque in the vessel. Although the patency of the blood vessel can be identified using standard diagnostic imaging techniques (MRI, CT and ultrasound) it has been shown that improved specificity for stroke prediction can be gained from identifying plaque features, such as intraplaque haemorrhage (IPH), that are associated with increased stroke risk^4,5^.

### 1.2 Quantitative Susceptibility Mapping

Quantitative susceptibility mapping (QSM) is an MRI method capable of spatially mapping the magnetic susceptibility of biological tissue^6–8^. Recently, a number of studies have applied QSM to carotid plaques and demonstrated marked improvement in IPH and calcification depiction compared to conventional multi-contrast MRI^9–13^ based on the inherent differences between the magnetic susceptibilities of these structures: calcifications are diamagnetic, whereas haemorrhages are paramagnetic^14^.

### 1.3 Arterial microstructure and QSM

Although QSM has proven promising for evaluating carotid plaques, specific disease driven sources of susceptibility changes have yet to be investigated in arterial vessels and plaques. Arterial tissue is predominantly composed of smooth muscle cells, elastin and collagen that are helically arranged to form the vessel microstructure^15^. The quantity, quality and organisation of this microstructure is finely tuned to maintain proper physiological function in healthy blood vessels^16^. Furthermore, arterial microstructure has been shown to become disrupted during the onset and progression of atherosclerosis and this changing microstructural arrangement may be indicative of mechanical stability, and therefore rupture risk, of advanced plaques^17^. As the magnetic susceptibility of a tissue is governed by its molecular makeup, QSM can provide insight into the microstructural composition of biological tissue. To date, QSM has been demonstrated to be sensitive to various tissue microstructural components such as myelin in brain^18,19^, collagen in liver^20,21^ and articular cartilage^22–24^, tubules in kidney^25^ and myofibers in heart^26^. As such, investigation of the sensitivity of QSM to arterial microstructure is warranted to determine if this is a source of susceptibility contrast that may provide an important biomarker of disease onset and progression in arteries.

### 1.4 Aim and summary

In this study we hypothesize that the magnetic susceptibility of arterial vessels is sensitive to changes in the vessel wall microstructure. To test this, a high resolution QSM protocol was applied to excised porcine carotid arteries subjected to a range of enzymatic digestion treatments^27^. To relate the magnetic susceptibility of porcine tissue to human arterial tissue, to accelerate potential clinical translation, the same QSM protocol was applied in *ex vivo* human carotid arteries. In both porcine and human vessels, detailed histological analysis was used to understand the underlying tissue microstructural composition and interpret the magnetic susceptibility of arterial vessels measured using QSM.

## 2. Methods

### 2.1 Sample preparation

#### 2.1.1 Porcine carotid artery (microstructural) models

Carotid arteries were harvested from 6-month-old, Large White pigs within three hours of slaughter. All vessels were cleaned of connective tissue and cryopreserved using a protocol to preserve the integrity of tissue microstructure during freezing^28^. In brief, cryopreservation was performed at a controlled rate of -1°C/min to -80°C in tissue freezing media composed of Gibco RPMI 1640 Medium (21875034, BioSciences), sucrose (S0389, Sigma) and the cryoprotectant dimethylsulfoxide (PIER20688, VWR International). The use of a cryoprotectant prevented microstructural damage caused by the formation of ice crystals^29^. In preparation for imaging, vessels were thawed at 37°C and rinsed in phosphate buffered saline (PBS) (P5493, Sigma).

To investigate the sensitivity of QSM to arterial tissue microstructure, four different vessel models were developed with distinct microstructural compositions using porcine carotid arteries. All vessels were cryopreserved prior to treatment and imaged directly after treatment and before fixation. The *native* vessel model refers to porcine carotid arteries that were not subjected to any treatment and acted as a control tissue for comparison with the following models.

The *collagen degraded* model was produced by incubating native porcine carotid arteries in 1000 U/ml purified collagenase (CLSPA, Worthington Biochemical Corporation) in magnesium chloride and calcium chloride supplemented PBS (D8662, Sigma) at 37°C for 28 hours on a rotator at 10 rotations per minute.

The *decellularised* model was produced using an established protocol as follows^30^: using a pressure of 100 mmHg, 0.1 M sodium hydroxide (S8045, Sigma) was perfused through native porcine carotid arteries via a peristaltic pump at 2 Hz for 15 hours, followed by 0.1 M sodium chloride (S3014, Sigma) for 32 hours. Vessels were then treated with 10 µl/ml DNAase (LS006343, Worthington Biochemical Corporation) and 2 µl/ml primicin (Ant-pm-2, InvivoGen) at 37°C for 19 hours.

The *elastin degraded* model was produced by incubating native porcine carotid arteries in 10 U/ml purified elastase (ESFF, Worthington Biochemical Corporation) with 0.35 mg/ml trypsin inhibitor (10109886001, Sigma) in Dulbecco’s Modified Eagle Medium, high glucose, GlutaMAX (61965026, BioSciences) at 37°C for 3.5 hours.

#### 2.1.2 Human carotid artery

Carotid arteries (including the carotid bifurcation and proximal sections of the common, internal and external carotid artery) were excised from five embalmed cadavers. One artery was obtained from each subject. The subjects (3 females and 2 males) ranged from 70 to 103 years in age (mean 81.6 ± 12.7 years). Cardiovascular disease was not implicated as the cause of death in any subjects. Vessels were cleaned of connective tissue and stored in PBS. To facilitate comparison of susceptibility values between human and porcine arteries, *fixed native* porcine carotid arteries were produced by immersing native porcine vessels in 4% formalin (HT501128, Sigma) for 7 days at 4°C. Prior to MR imaging all samples were washed and placed in fresh PBS.

### 2.2 MR imaging

#### 2.2.1 Vessel positioning

All vessels were positioned using 3D-printed holders composed of polylactic acid and placed in 50-ml falcon tubes (Supporting Information **Figure S1**) in which samples were immersed in fresh PBS prior to imaging at room temperature. PBS was chosen as the “embedding” material as it has previously been identified as providing a stable experimental set-up facilitating good image quality for QSM of post-mortem brain specimens^31^. For the porcine vessels (*native, decellularised, collagen degraded, elastin degraded* and *fixed native*), six vessels (n=6) of each model were produced and holders were designed to secure six vessels in a single 50 ml falcon tube. For human vessels, holders were designed to hold a single specimen.

#### 2.2.2 Image acquisition

A small bore (30 cm) horizontal 7T Bruker BioSpec 70/30 USR system (Bruker, Ettlingen Germany) equipped with a receive-only 8-channel surface array coil, birdcage design transmit coil, shielded gradients (maximum strength 770 mT/m) and Paravision 6 software was used for all imaging. For each session, a test tube containing the vessels of interest was placed securely in the cradle of the 8-channel surface array coil. A total of ten scan sessions were performed (five sessions for porcine vessels and five sessions for human vessels).

For QSM, data were acquired using a 3D multi-echo gradient echo (ME-GRE) sequence with the following parameters: TEs = 5, 13.1, 21.2 and 29.3 ms with monopolar readout gradients, TR = 150 ms, flip angle = 30°, bandwidth = 34722 Hz and averages = 2. The readout direction was oriented along the long axis of the tube. An isotropic voxel resolution of 0.117 x 0.117 x 0.117 mm^3^ was achieved using a field-of-view of 30 mm x 30 mm x 30 mm and a 256 x 256 x 256 matrix size. Total scan time for this sequence was 5 hours 27 minutes.

### 2.3 Image Processing

For all samples, the multi-channel ME-GRE data was coil-combined. Coil-combined magnitude images were calculated using the root sum of squares of the channels^32^ and coil-combined phase images were produced using the phase difference approach^33^.

#### 2.3.1 QSM pipeline in porcine vessels

The processing pipeline used to produce quantitative susceptibility and relaxometry maps from the imaging data acquired in porcine vessels is summarised in **Figure 1**. R_2_* maps were calculated using the Auto-Regression on Linear Operations algorithm (ARLO)^34^ applied to the coil-combined ME-GRE magnitude data. To aid masking, an echo-combined magnitude image was calculated using the root sum of squares of all echoes. For QSM, an initial mask was created by thresholding the echo-combined magnitude image with the threshold set to include all vessels and PBS but to exclude the 3D-printed vessel holder and air outside the tube. The mask was manually refined to exclude air bubbles using the echo-combined magnitude and R_2_* map as references. Non-linear field fitting^35,36^ was used to estimate field maps from the complex ME-GRE data and remaining wraps were removed using Laplacian phase unwrapping^37^. This approach (non-linear field fitting followed by Laplacian phase unwrapping) provided a computationally efficient and robust approach to calculating unwrapped total field maps and has previously been applied to investigate tissue susceptibility in *ex vivo* articular cartilage^22^. Alternative approaches, such as linear field fitting, require prior unwrapping of individual echoes, which would lead to long computation times particularly for the large matrix sizes acquired here (256 x 256 x 256) whereas the approach used in this study requires unwrapping of only a single image. The local field map was calculated using the projection onto dipole fields (PDF) method and the unwrapped field map and magnitude mask as input^38,39^. PDF was chosen as it has been shown to perform well in comparison to alternative methods^40^. A susceptibility map was calculated from the local field map using an iterative Tikhonov method^7,41,42^ with correction for susceptibility underestimation^43^. The same regularisation parameter (α) was used for all samples and was chosen by performing L-curve optimisation^44^ in all five porcine datasets and calculating the mean of the individually optimised parameters. Alternative susceptibility calculation methods (TKD^45^, direct Tikhonov^7,41,46^ and MEDI^47^) were tested and found not to influence the trends and final conclusions reported for porcine vessels (see Supporting Information **Figures S9** and **S10**).

**Figure 1.**
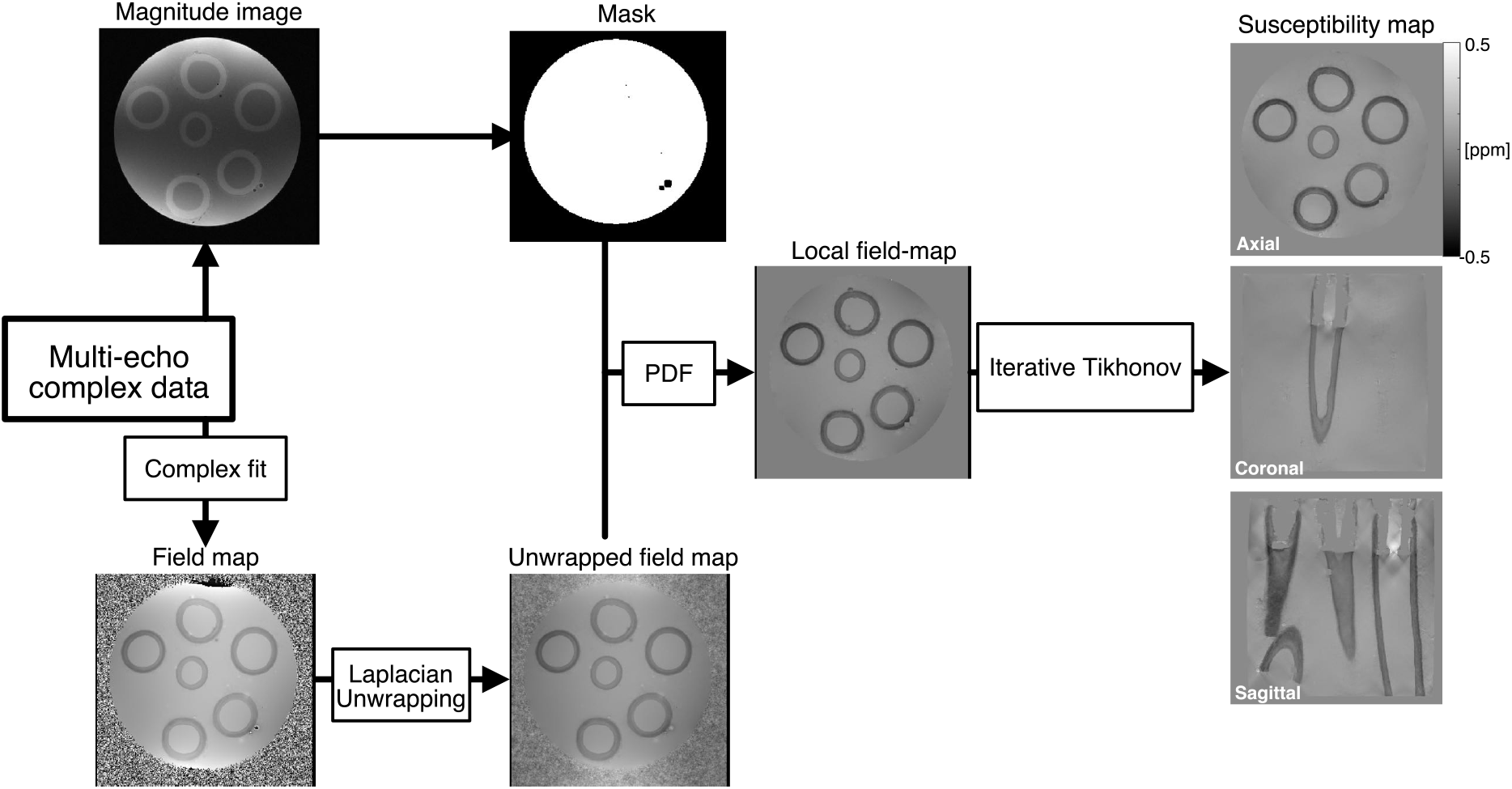
Outline of QSM pipeline for porcine carotid artery tissue models.

#### 2.3.2 QSM pipeline in human vessels

All five human vessels scanned in this study had regions of advanced atherosclerotic disease close to the bifurcation (see **Figure 7** yellow arrows). These heavily diseased regions contained structures with little or no signal that caused significant streaking in the final susceptibility maps likely attributable to the presence of calcification or haemorrhage. To investigate the impact of these low signal-to-noise-ratio (SNR) regions on the susceptibility measurements made in regions of the common carotid unaffected by disease/plaque, two different masking procedures were compared for QSM (**Figure 2**).

**Figure 2.**
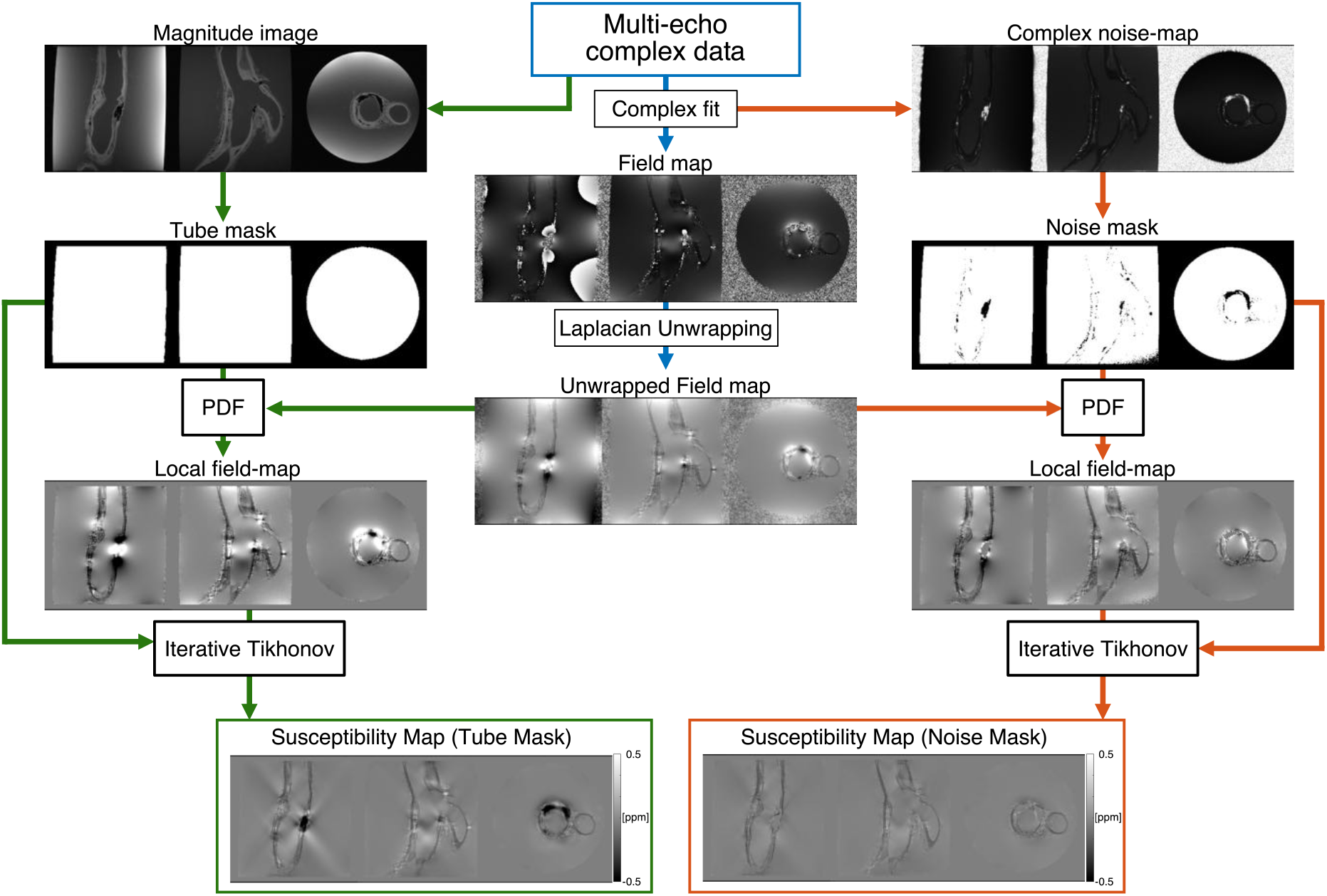
Outline of QSM pipeline for human carotid specimens. Two different masks were compared to investigate the impact of low SNR, diseased regions on the susceptibility measured in nearby healthy regions of the common carotid artery. The noise mask pipeline (orange) excluded regions of low SNR from the background field removal and susceptibility calculation steps using a mask that was calculated by thresholding the inverse of the residual noise map produced from the non-linear fit of the complex data. This was compared to a tube mask pipeline (green) that used a simple tube mask that was manually defined on the echo-combined magnitude image. This mask included all the material present in the tube and the high-noise regions.

**Figure 3.**
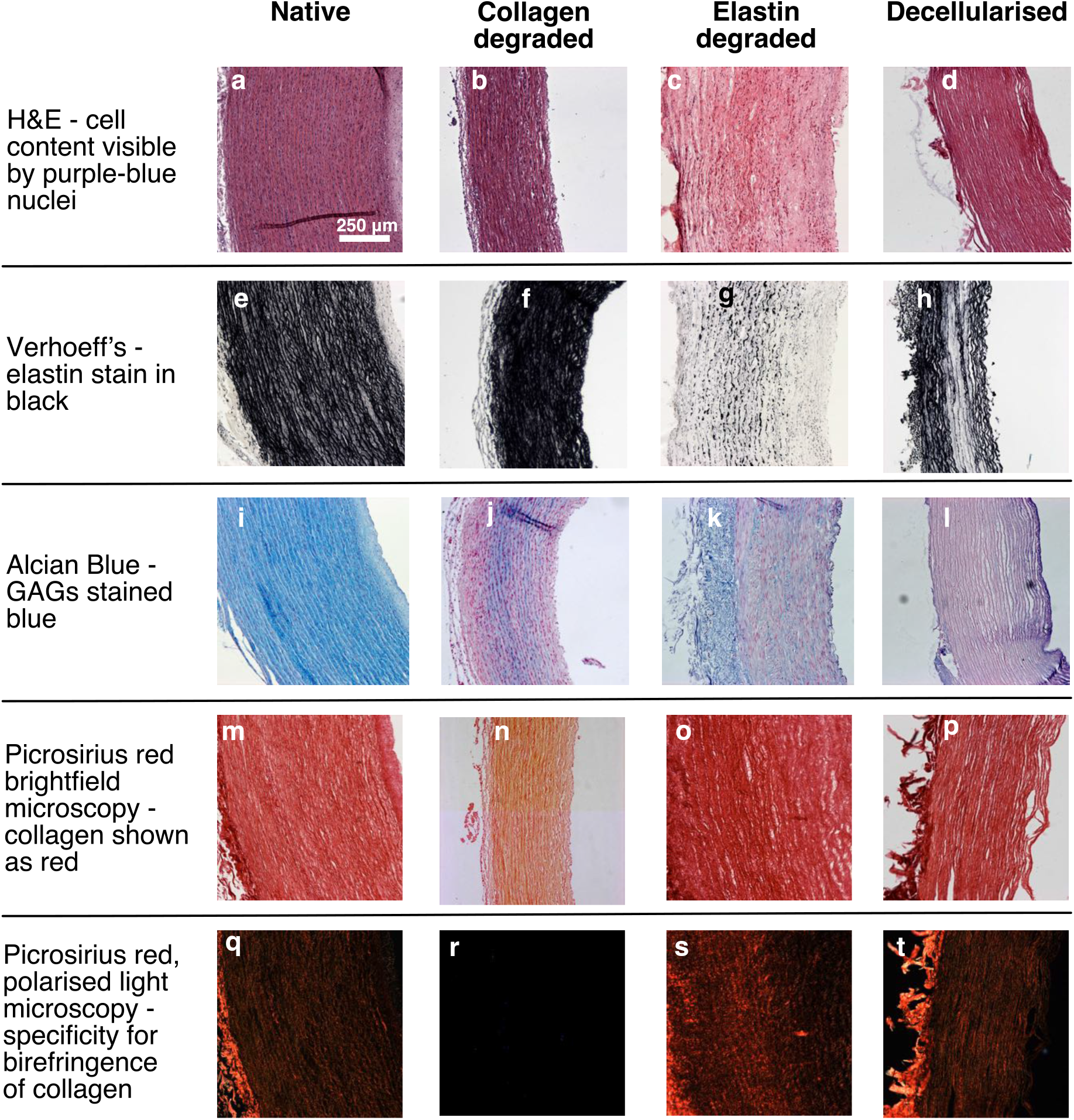
Histological validation of porcine carotid artery tissue models. Enzymatic digestion treatments were used to selectively remove components of the arterial microstructure creating four different tissue models – native, collagen degraded, elastin degraded and decellularised porcine carotid artery

##### 2.3.2.1 Comparison of masking procedures for human vessels

Firstly, a *tube mask* was generated for each human vessel that *included* heavily diseased low SNR regions. This mask was generated by thresholding the echo-combined magnitude image to exclude the 3D-printed vessel holder and air outside the tube and include all other contents of the tube, i.e. PBS and the whole vessel (including diseased regions). No manual refinement step was implemented here due to the difficulty in distinguishing diseased regions from air bubbles.

Secondly, a *noise mask* was generated for each human vessel that *excluded* heavily diseased low SNR regions. High noise regions in areas of advanced atherosclerotic disease were identified by thresholding the inverse noise map (calculated during non-linear field fitting^36^ using noise propagation^46^) at one third of the mean voxel value contained within the *tube mask*. To generate the final *noise mask*, the regions of high noise identified by the inverse noise map were removed from the *tube mask* (see **Figures 2** and **8**).

##### 2.3.2.2 QSM calculation in human vessels

Masking procedures were compared for final susceptibility map calculation through comparison of *tube mask* and *noise mask* QSM pipelines. For both pipelines, field maps were generated using non-linear field fitting^35,36^ and unwrapped using Laplacian phase unwrapping^37^. At this point, the pipelines diverge as background field removal and susceptibility calculation steps require masks identifying the region of interest. Local field maps were calculated with the PDF method^34^ using the unwrapped field map and a region of interest mask as input i.e. *tube mask* for the *tube mask* pipeline and *noise mask* for the *noise mask* pipeline. Using the relevant local field map and mask as inputs, susceptibility maps were calculated for the *tube mask* and *noise mask* pipelines using the iterative Tikhonov approach as described for the porcine arteries. The same regularisation parameter (α) was used for all samples and was chosen by performing L-curve optimisation^44^ for both *tube mask* and *noise mask* pipelines in all five human datasets and calculating the mean of the individually optimised parameters. Similar to the porcine vessels, alternative susceptibility calculation methods (TKD, direct Tikhonov and MEDI) were tested and found not to influence the trends and final conclusions reported for human vessels (see Supporting Information **Figure S10**).

### 2.4 Histological Analysis

To allow histological validation, the model tissues (*native, decellularised, collagen degraded* and *elastin degraded* vessel models) were fixed immediately after MR scanning for histological processing by immersing vessels in 4% formalin for seven days at 4°C. This was unnecessary for *fixed native* porcine and human vessels as fixation was performed prior to scanning. Stepwise dehydration was performed on the fixed tissue in ethanol to xylene and samples were then embedded in paraffin wax and sectioned into 8 µm thick slices prior to staining.

For porcine and human vessels, Haematoxylin and Eosin (H&E), Picrosirius red, Verhoeff’s elastin and Alcian blue staining were performed to identify the presence of smooth muscle cells, collagen, elastin and glycosaminoglycans (GAGs) respectively^27^. Additionally, Alizarin Red staining was performed on human vessels to identify the presence of calcium. Histological imaging was performed using an Olympus BX41 microscope with Ocular V2.0 software for the porcine models and an Aperio CS2 microscope with ImageScope software V12.3 for the human arteries. Bright-field microscopy was performed for all stains with additional polarised light microscopy (PLM) on Picrosirius red to infer the orientation of collagen fibres.

For human vessels, alignment of histology to QSM was performed as follows; using anatomical features as landmarks, the first echo of the magnitude combined ME-GRE image was used to guide the identification of tissue sections in the common carotid for histological analysis. The location of the selected region on the ME-GRE image was noted and used to guide the manual alignment of the digitised whole-mount histology image to the relevant MRI slice. The final alignment of histology to MRI was refined via manual rotation and scaling of the histology image (see **Figure 4** and Supporting Information **Figures S2-S6)**

**Figure 4.**
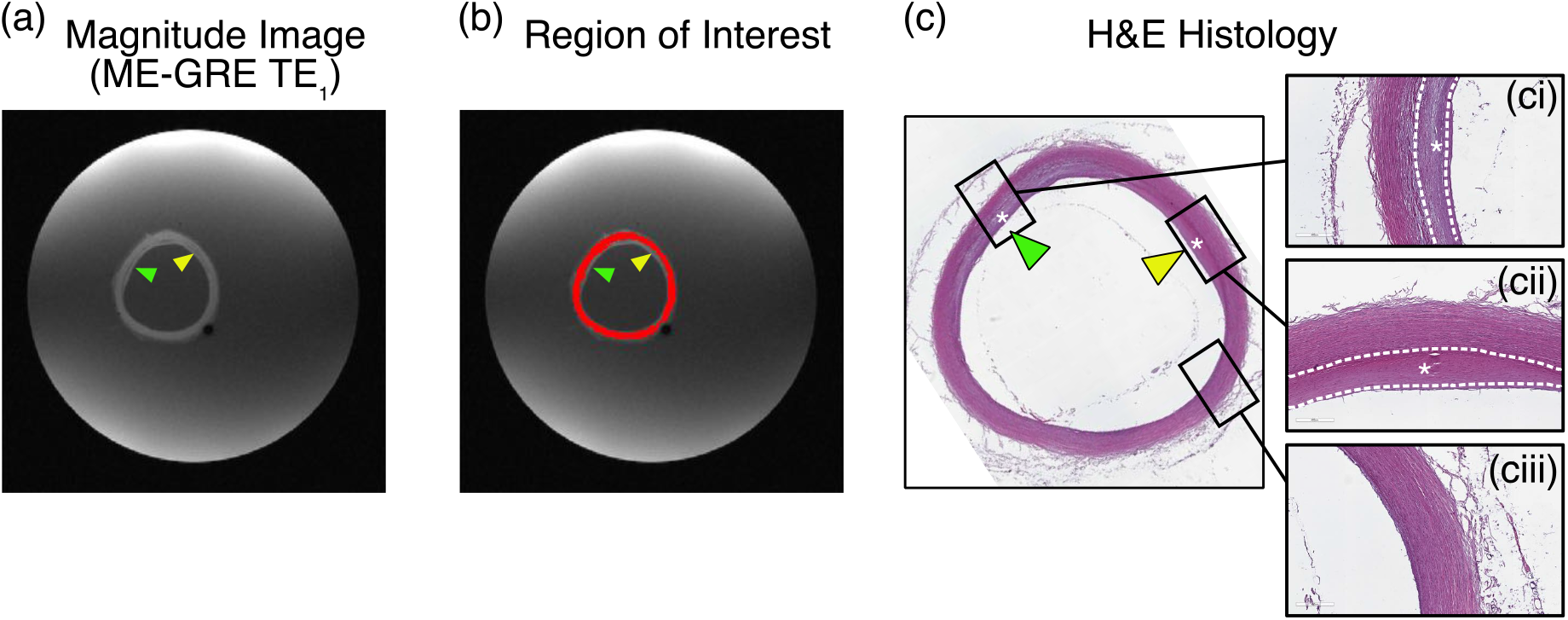
Example of MRI ROI definition in normal appearing human common carotid. ME-GRE images (a) with ROI overlay (b) are displayed alongside H&E histology (c). H&E histology identifies regions of intimal thickening (ci) and (cii) and healthy appearing vessel microstructure (ciii). Abnormal regions are excluded from ROI definition on the magnitude images (green and yellow arrows (b)). Detailed comparison of MRI ROIs and histological analysis for each vessel are shown in Supporting Information **Figures S2-S6**

### 2.5 Data Analysis

#### 2.5.1 Regions of interest

For both human and porcine tissue, regions of interest (ROIs) were manually defined for each vessel using the echo-combined MRI magnitude image. Care was taken to position ROIs within arterial tissue and avoid regions of partial volume near the edges of each vessel. This was facilitated by the high resolution of the ME-GRE data. ROIs in human vessels were limited to normal appearing arterial tissue in the common carotid artery. This was achieved by cross referencing the MRI defined ROI with histology and excluding regions from the final ROI that showed abnormal smooth muscle cell, collagen, elastin, GAG or calcium content as identified by histology and defined by *Stary et al*.^48^ (see **Figure 4** and Supporting Information **Figures S2-S6**). Using the echo-combined magnitude image, ROIs were manually defined for PBS in each sample. Following ROI definition, ROI mean susceptibility values were extracted for each vessel and for PBS in each sample. To compare across samples, vessel susceptibility was referenced to PBS^49^. This was achieved by subtracting the mean PBS susceptibility from the mean vessel susceptibility in the same sample.

### 2.5.2 Statistical methods

For measurements made in porcine vessels, the null hypothesis of no susceptibility difference between tissue models (*native, decellularised, collagen degraded, elastin degraded* and *fixed native*) was tested using a one-way ANOVA. If the null hypothesis was rejected, post hoc pairwise comparisons were performed using the Tukey-Kramer method. To compare the susceptibility measurements made using *tube mask* and *noise mask* pipelines in human common carotid arteries, the intraclass correlation coefficient (ICC)^50^ was calculated. For comparison of susceptibility measurements made in human common carotid arteries with those in *fixed native* porcine arteries, the null hypothesis of no susceptibility difference between groups (human common_tube mask_, human common_noise mask_ and *fixed native* porcine) was tested using a one-way ANOVA. If the null hypothesis was rejected, post hoc pairwise comparisons were performed using the Tukey-Kramer method.

## 3. Results

### 3.1 Histology

#### 3.1.1 Histological validation of tissue models

Figure 3 presents histological validation of the vessel models where enzymatic digestive treatments were used to selectively remove smooth muscle cells, elastin and collagen in selected groups of porcine carotid arteries. H&E staining confirms the absence of smooth muscle cells in the *decellularised* vessels (**Figure 3 d**), Verhoeff’s elastin stain confirms the degradation of elastin in the *elastin degraded* vessels (**Figure 3 g**) and Picrosirius red verifies the removal of collagen in the *collagen degraded* vessels (**Figure 3 n** and **r**). Furthermore, H&E staining (**Figure 3 a-d**) verifies that smooth muscle cell content remained intact in the non-decellularised vessels, Verhoeff’s elastin staining (**Figure 3 e-h**) verifies that elastin content was maintained in the non-elastin degraded vessels and Picrosirius red staining with bright field microscopy (**Figure 3 m-p**) and PLM (**Figure 3 q-t**) verifies the preservation of collagen content and orientation in the non-collagen degraded vessels. Alcian blue staining (**Figure 3 i-l**) showed a decrease in GAG content across all the degradation models when compared with native tissue. This may be explained by the leaching of GAGs out of the tissue in order to maintain osmotic balance due to the presence of PBS. Although this hasn’t been observed in arterial tissue it has been observed in intervertebral disc and articular cartilage^51^.

#### 3.1.2 Histological validation of ROIs in cadaver vessels

Figure 4 shows example H&E in human common carotid artery for a representative cadaver specimen imaged in this study. Supporting Information **Figures S2-S6** display histological analysis and ROI definition for each human vessel. Histology was used to guide the definition of MRI ROIs in “normal” appearing common carotid tissue by avoiding regions of obvious abnormality or disease defined as a disruption to cell, collagen, GAG, elastin or calcium content.

### 3.2 MRI

#### 3.2.1 QSM of tissue models

Figure 5 presents susceptibility and R_2_* relaxometry maps for each of the vessel models. Susceptibility maps produced using the iterative Tikhonov method are presented in three different viewing planes (sagittal, coronal and axial in the context of the tube which was placed with its long axis parallel to the orientation of B_0_) for qualitative comparison of image quality. The regularisation parameter used for susceptibility map calculation in porcine vessels was α=0.02 as determined by taking the mean of the L-curve optimisation^44^ across the five samples (Supporting Information **Figures S7**).

Qualitative differences are apparent when comparing the susceptibility maps of different vessel models. The susceptibility of *native* (**Figure 5 a**) vessels appear the most diamagnetic with the susceptibility of *decellularised* (**Figure 5 b**) and *collagen degraded* (**Figure 5 c**) vessels appearing elevated in comparison while maintaining contrast with the background fluid. The susceptibility of the *elastin degraded* vessels (**Figure 5 d**) appear the most elevated with almost no susceptibility difference with the surrounding fluid. Qualitatively, R_2_* relaxometry maps demonstrated similar but inverse trends to the susceptibility maps with R_2_* of *native* vessels (**Figure 5 e**) appearing highest and R_2_* of *elastin degraded* vessels (**Figure 5 h**) appearing lowest and similar to that of surrounding background fluid.

**Figure 5.**
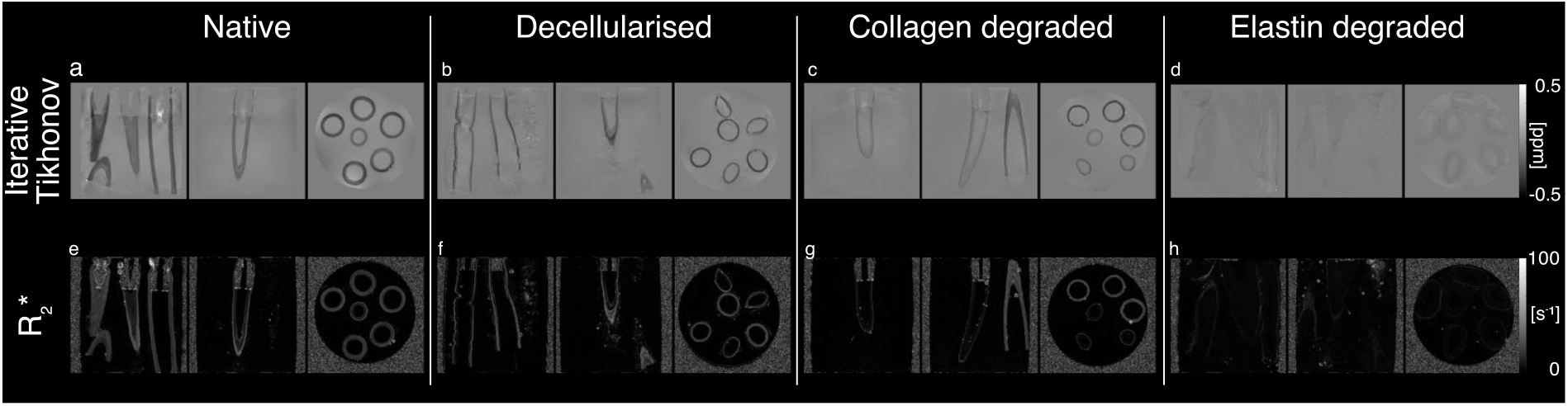
Susceptibility maps produced for the different porcine carotid tissue models (native, decellularised, collagen degraded and elastin degraded) using the iterative Tikhonov approach (a-d) alongside maps of R_2_* (e-h). Equivalent susceptibility maps calculated with alternative susceptibility calculation methods (TKD, direct Tikhonov and MEDI) are compared in Supporting Information **Figures S8**

As a quantitative comparison, **Figure 6** shows boxplots of the susceptibility and R_2_* values measured in each vessel (n=6 per model) and grouped by tissue model. The mean (± standard deviation) susceptibility of PBS measured across the five samples was 0.0219 ± 0.0121 ppm. Susceptibility and R_2_* measurements in *fixed native* porcine carotid artery (n=6) are presented for later comparison with human vessels. The baseline magnetic susceptibility of *native* porcine arteries was found to be diamagnetic with a mean value of −0.1820 ppm and vessel susceptibilities ranged from −0.2346 ppm to -0.0092 ppm across all tissue models. The mean susceptibility of the tissue models (𝒳_elastin degraded_ = −0.0163 ppm; 𝒳_collagen degraded_ = −0.1158 ppm; 𝒳_decellularised_ = −0.1379 ppm; 𝒳_fixed native_ = −0.2199 ppm) exhibited the following trend: 𝒳_elastin degraded_ > 𝒳_collagen degraded_ > 𝒳_decellularised_ > 𝒳_native_ > 𝒳_fixed native_. Significant differences were detected between the susceptibility of the vessel models (ANOVA, p<0.001). Post hoc pairwise comparisons revealed the susceptibility of *collagen degraded* vessels to be significantly higher than *native vessels* (p<0.01), the susceptibility of *elastin degraded* vessels to be significantly higher than all other vessel groups (p<0.001) and the susceptibility of *fixed native* vessels to be significantly lower than both *decellularised* (p<0.01) and *collagen degraded* vessels (p<0.001). Comparing vessel-wise measurements of R_2_*, the null hypothesis was rejected (p<0.001) indicating differences between groups. Post hoc testing revealed significantly lower R_2_* in *elastin degraded* vessels compared to all other vessel models (p<0.001).

**Figure 6.**
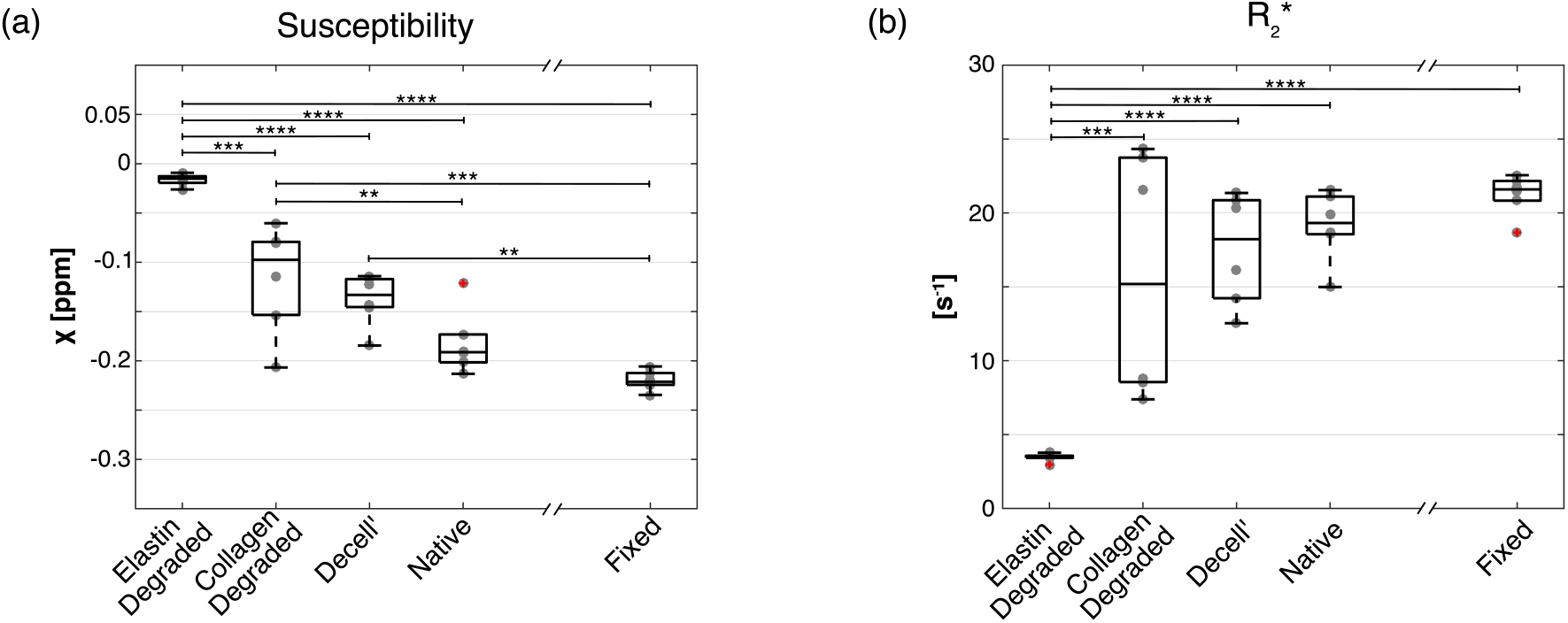
Boxplots comparing measurements extracted from vessels (n = 6) of each tissue model for QSM (a) and R_2_* maps (b). Measurements compared using ANOVA and post-hoc test (* p < 0.05, ** p < 0.01, *** p < 0.001, **** p < 0.0001). Equivalent measurements made using alternative susceptibility calculation methods are presented in Supporting Information **Figure S9**

#### 3.2.2 QSM of human common carotids

In the human carotid arteries, regions of advanced disease were seen close to the bifurcation in all five vessels scanned in this study (**Figure 7** yellow arrows). These heavily diseased regions contained structures with little or no signal – likely attributable to the presence of calcification or haemorrhage. For a representative vessel, susceptibility maps are presented for *tube mask* and *noise mask* pipelines, with the region where the ROI was defined, indicated in red (**Figure 8**). It can be seen that human carotid vessels exhibited pronounced streaking artefacts attributed to the inclusion of low SNR regions in the QSM calculation (*tube mask*). Streaking was visibly reduced in the susceptibility maps produced using the *noise mask* pipeline (**Figure 8**). The optimal regularisation parameter was determined as α=0.02 using L-curve optimisation, remaining the same as that used for the porcine vessels (Supporting Information **Figures S7**).

**Figure 7.**
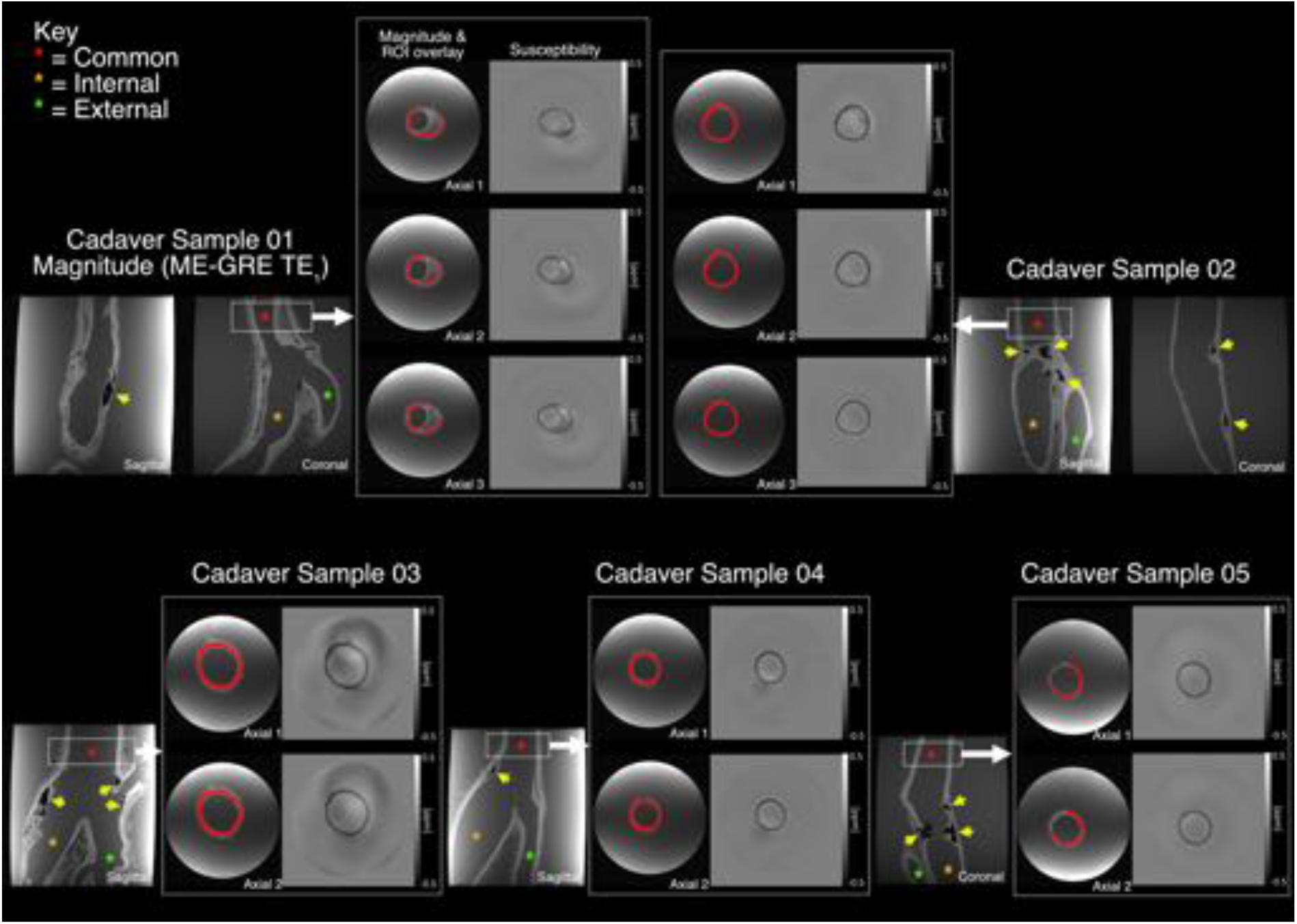
Measurements in human cadaver samples were confined to “normal” common carotid artery tissue with no areas of atherosclerotic disease. The location of the common carotid ROI is displayed on first-echo magnitude images from the ME-GRE acquisition. Common (red asterisk), internal (yellow asterisk) and external (green asterisk) carotid arteries can be seen in the sagittal / coronal magnitude images. Representative QSM axial images in the common carotid ROI are presented alongside first-echo magnitude axial images with red ROI overlaid

**Figure 8.**
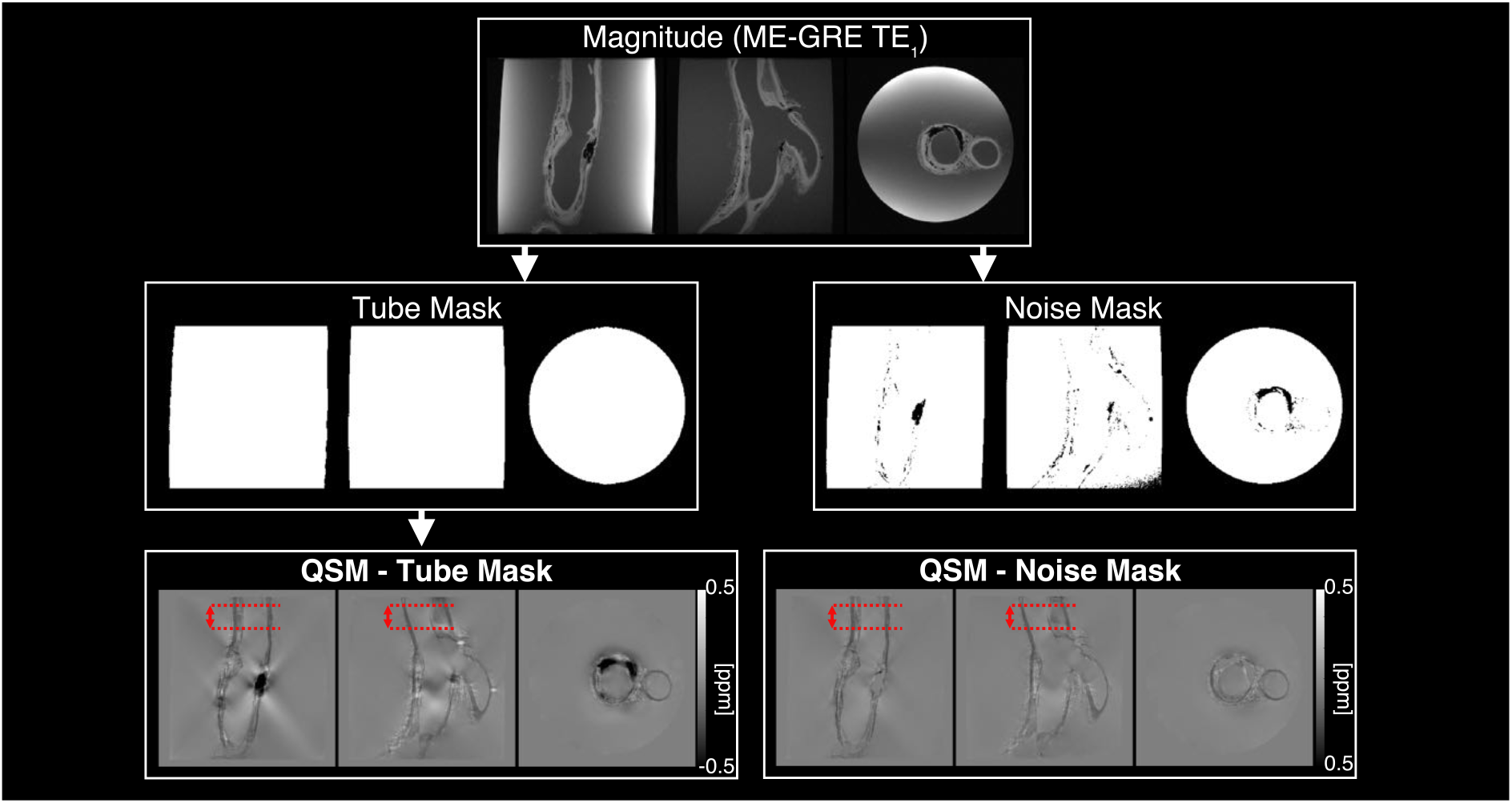
Visual comparison of QSM pipelines including and excluding high-noise regions (*tube mask* and *noise mask*) in a representative human cadaver carotid artery. The common carotid ROI was defined in “normal” appearing tissue between the red dotted lines

Figure 9 shows susceptibility measurements in human common carotid arteries. The mean susceptibility of human common carotid was −0.1898 ± 0.0253 ppm using the *noise mask* pipeline and −0.2007 ± 0.0183 ppm using the *tube mask* pipeline. This is in comparison to the *fixed native* porcine carotid arteries which had a mean susceptibility of −0.2199 ± 0.0100 ppm. **Figure 9 a** directly compares common carotid susceptibility measurements between the *noise mask* and *tube mask* pipelines with an ICC value of 0.88 (p<0.05) being obtained between the pipelines. **Figure 9 b** displays box-plots of the mean susceptibility in each vessel using the *noise mask* and *tube mask* pipelines compared with those in *fixed native* porcine carotid arteries. The null hypothesis was not rejected (p>0.05), however the p-value of 0.050 was close to the significance level of 0.05.

**Figure 9.**
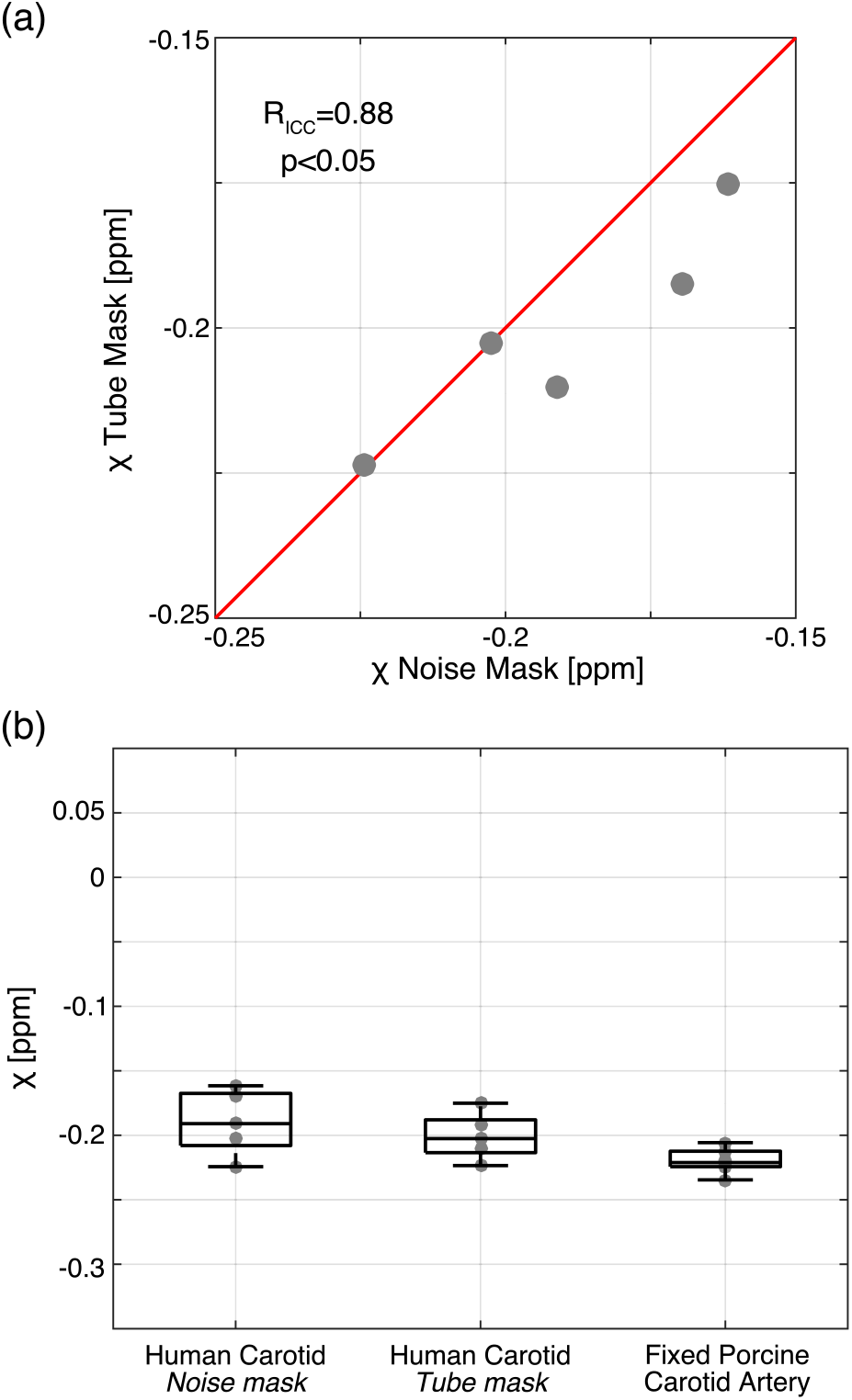
Magnetic susceptibility of human common carotid. The effect of masking out noisy regions on mean susceptibility values in the cadaver common carotid ROI are shown in **(a)**. Agreement between different masking pipelines was excellent using the following criteria - Poor < 0.4, Fair - 0.41–0.59, Good - 0.60– 0.74, Excellent > 0.75^52^. Boxplots comparing susceptibility measurements in cadaver common carotid arteries with those in fixed porcine carotid arteries is shown in **(b)**. Susceptibility values were compared using ANOVA and no significant difference was seen between groups (p=0.0501). Equivalent measurements made using alternative susceptibility calculation methods are presented in Supporting Information **Figure S10**

## 4. Discussion

### 4.1 Main findings

In this study we demonstrate for the first time the sensitivity of magnetic susceptibility, measured using QSM, to the microstructural composition of arterial tissue. Vessels with different microstructural compositions were generated by applying different enzymatic digestion treatments to *ex vivo* porcine carotid arteries. For each model (*decellularised, collagen degraded* and *elastin degraded*), six vessels were imaged using a high resolution QSM protocol and compared to untreated (*native*) and *fixed native* porcine carotid arteries (**Figure 5**). The microstructural composition of each vessel model was validated using histology (**Figure 3**) and statistically significant differences were found between the group mean susceptibility of porcine vessels with different microstructural compositions (**Figure 6**). Post hoc statistical testing revealed significantly higher susceptibility in *collagen degraded* vessels compared to *native* vessels, while no such difference was present in equivalent R_2_* relaxometry measurements. Significantly lower susceptibility was measured in *fixed native* vessels compared to both *decellularised* and *collagen degraded* vessels with no such statistical differences existing in the equivalent R_2_* measurements. This suggests that QSM offers improved sensitivity to the microstructural composition of arterial vessels, in particular collagen, when compared to equivalent R_2_* measurements.

To provide a comparison with human tissue, susceptibility was measured in *ex vivo* human common carotid arteries using the same high resolution QSM pipeline (**Figure 7**). Regions of advanced disease were present in these vessels leading to streaking artefacts in the susceptibility maps. Exclusion of these low SNR regions resulted in a qualitative improvement in the susceptibility maps (**Figure 8**). However, using ICC “excellent” agreement^52^ was found between human common carotid measurements extracted from susceptibility maps that were calculated using pipelines which included and excluded the low SNR diseased regions (**Figure 9 a**). One explanation for this agreement is that the common carotid is positioned far enough away from the streaks (originating in the bifurcation) that they have very little effect on the mean susceptibility measured in that region. An example of this can be seen in **Figure 8**. Statistical testing revealed no significant differences between magnetic susceptibilities measured in fixed human common carotid arteries and fixed porcine vessels (**Figure 9 b**).

### 4.2 Susceptibility and arterial microstructure

Experimental results from porcine vessel models (**Figure 6**) suggest tissue susceptibility, measured using QSM, is sensitive to the microstructural composition of arterial vessels. The baseline magnetic susceptibility of native porcine arteries was found to be diamagnetic with a mean value of −0.1820 ppm and agrees well with the value of −0.25 ± 0.14 ppm reported for popliteal artery wall *in vivo*^24^.

#### 4.2.1 Collagen

Compared to *native* vessels, a significantly higher susceptibility was measured in *collagen degraded* vessels (Δ𝒳 = 0.0662 ppm) while no such significant difference was apparent in the equivalent R_2_* relaxometry measurements. This agrees well with studies of collagen susceptibility in articular cartilage^24,53^ and liver fibrosis^21^ which observed collagen as strongly diamagnetic. Although articular cartilage degradation associated with collagen loss has been shown to cause changes in measured susceptibility *in vivo*^54^, *ex vivo* studies of articular cartilage failed to detect differences in the susceptibility of collagen degraded samples^22^. The differences in the results found in *ex vivo* collagen degraded articular cartilage and arterial vessels may well be explained by differences in experimental set-up, susceptibility reference material and enzymatic digestion treatments. It is important to note that collagen was completely removed from the samples imaged in this study (**Figure 3 n & r**) and PBS was used as a consistent susceptibility reference between samples.

#### 4.2.2 Elastin

Significant differences were seen between the measured susceptibility of *elastin degraded* vessels and all other vessel models including *native* and *fixed native* vessels. The *elastin degraded* group also demonstrated the largest deviation in measured susceptibility from *native* vessels. However, as evidenced histologically by H&E staining (**Figure 3 a** & **c**) the removal of elastin results in a less compact microstructural arrangement of tissue, when compared to *native* vessel histology. This suggests that the removal of elastin results in an increased extracellular space, allowing the penetration of surrounding PBS into the tissue microstructure. This is supported by the visible increase in size of the vessels and the distinct lack of contrast seen between *elastin degraded* vessels and the surrounding PBS in QSM (**Figure 5 d**). Therefore, it is difficult to conclusively assess the contribution of elastin to vessel susceptibility from the tissue model presented here.

#### 4.2.3 Smooth muscle cells

Considerable overlap is seen between the measured susceptibilities of *decellularised* and *native* vessels. The slightly higher group mean susceptibility of *decellularised* vessels was not significantly different from that of *native* vessels (**Figure 6**).

The results from all these porcine artery models suggest that QSM is most sensitive to detecting changes in arterial collagen. Further work is required to investigate the integrity of the elastin model and the resulting abolition, following elastin degradation, of the diamagnetic susceptibility found in native porcine arteries. An alternative approach to characterising the magnetic susceptibility of elastin is to use 3D printed tissue scaffolds where the density of elastin can be tightly controlled^55^.

#### 4.2.4 Susceptibility anisotropy

Although this study has demonstrated the sensitivity of QSM to arterial microstructure, specific components, such as collagen, are known to possess B_0_ orientation-dependent magnetic susceptibility^21–23^. In this study, all vessels were imaged at the same orientation to the main magnetic field (long axis of the vessel parallel to B_0_). As arterial collagen and elastin fibres are circumferentially arranged, these fibres will be oriented at 90° to B_0_, facilitating consistent comparison of measured susceptibility within and between vessels. As QSM assumes isotropic susceptibility, further work is required to assess the susceptibility anisotropy of arterial tissue and the anisotropic contributions of its microstructural components, in particular collagen^53^.

### 4.3 Human artery

#### 4.3.1 Comparison of human and porcine vessels

From **Figure 9 b**, variability of vessel susceptibility across the group is visibly lower for porcine vessels compared to the human samples. This may be partially attributed to the low variability in the age of the pigs compared to the human subjects, but also to differences between fixation and embalming, respectively. Porcine vessels were imaged directly after fixation whereas time from embalming was not controlled for in human vessels. *Evia et al*. reported no systematic change in magnetic susceptibility of fixed post-mortem brain over a period of 6 weeks^56^ but changes in susceptibility outside this timeframe may be possible. Although specific information regarding time from embalming to scanning is not available for the human vessels, imaging of all vessels included in this study occurred after this 6-week time frame. The variability in susceptibility seen between human vessels could also be due to biological variability (e.g. age, sex, body size, disease status), as well as its effect on the embalming protocol.

Despite this variability there was no statistically significant difference measured between the magnetic susceptibility of human and porcine arterial tissue. This suggests that the sensitivity of magnetic susceptibility to microstructural composition, demonstrated in porcine arterial tissue, is also likely to be found in human arterial tissue. This provides a promising springboard for studies seeking to translate QSM for use in characterising human carotid arteries and disease.

#### 4.3.2 Fixed porcine tissue

The group mean susceptibility was slightly lower in fixed tissue (𝒳 = −0.2199 ppm) compared to native vessels (𝒳 = −0.1820 ppm), although this difference was not statistically significant (**Figure 6**). As noted by *Wei et al*.^21^, fixation alters tissue microstructure through crosslinking of proteins and differences in measured susceptibility are not surprising. Differences in susceptibility between *in vivo* and fixed *ex vivo* mouse brain have been reported^57^ and it has been noted that changes in tissue relaxation times that accompany fixation could impact QSM measurements of susceptibility^26^.

### 4.4 Conclusion, impact and future work

In diseased arterial tissue, key components of the tissue microstructure, such as collagen fibres, smooth muscle cells and elastic lamina become disrupted, impairing vessel function^58^. A number of recent studies have demonstrated the application of QSM for imaging carotid plaque *in vivo*, highlighting that many technical challenges associated with imaging carotid arteries using QSM in a clinical setting can be overcome^9–13^. These studies have demonstrated marked improvement in depiction of IPH and calcification *in vivo* but have largely ignored differences in regional plaque susceptibility that may be driven by compositional variations in the microstructure of fibrotic plaque tissue. Results from this study highlight that QSM is sensitive to the microstructural composition of arterial tissue and, with further development, has the potential to offer unique insight into the onset and progression of carotid atherosclerosis. Such characterisation of carotid plaques has the potential to improve the assessment of stroke risk using MRI^59^ and could complement existing MRI methods capable of detecting downstream haemodynamic alterations^60,61^. Future work will focus on QSM of diseased arterial tissue *ex vivo*, using the insights from this study as a basis to fully characterise the susceptibility contributions from IPH, calcifications, lipid and tissue microstructure in heterogenous atherosclerotic plaques.

## Acknowledgements

The authors would like to thank and acknowledge the Department of Anatomy, Royal College of Surgeons in Ireland (Professor Clive Lee and Bob Dalchan) for supporting this work

## Data Availability Statement

The imaging data for this study is available from the corresponding author on reasonable request. Tools used for QSM calculation are openly available through the MEDI toolbox (http://weill.cornell.edu/mri/pages/qsm.html) and UCL’s XIP repository (https://xip.uclb.com/i/software/mri_qsm_tkd.html)

## Supporting Information

**Figure S1:**
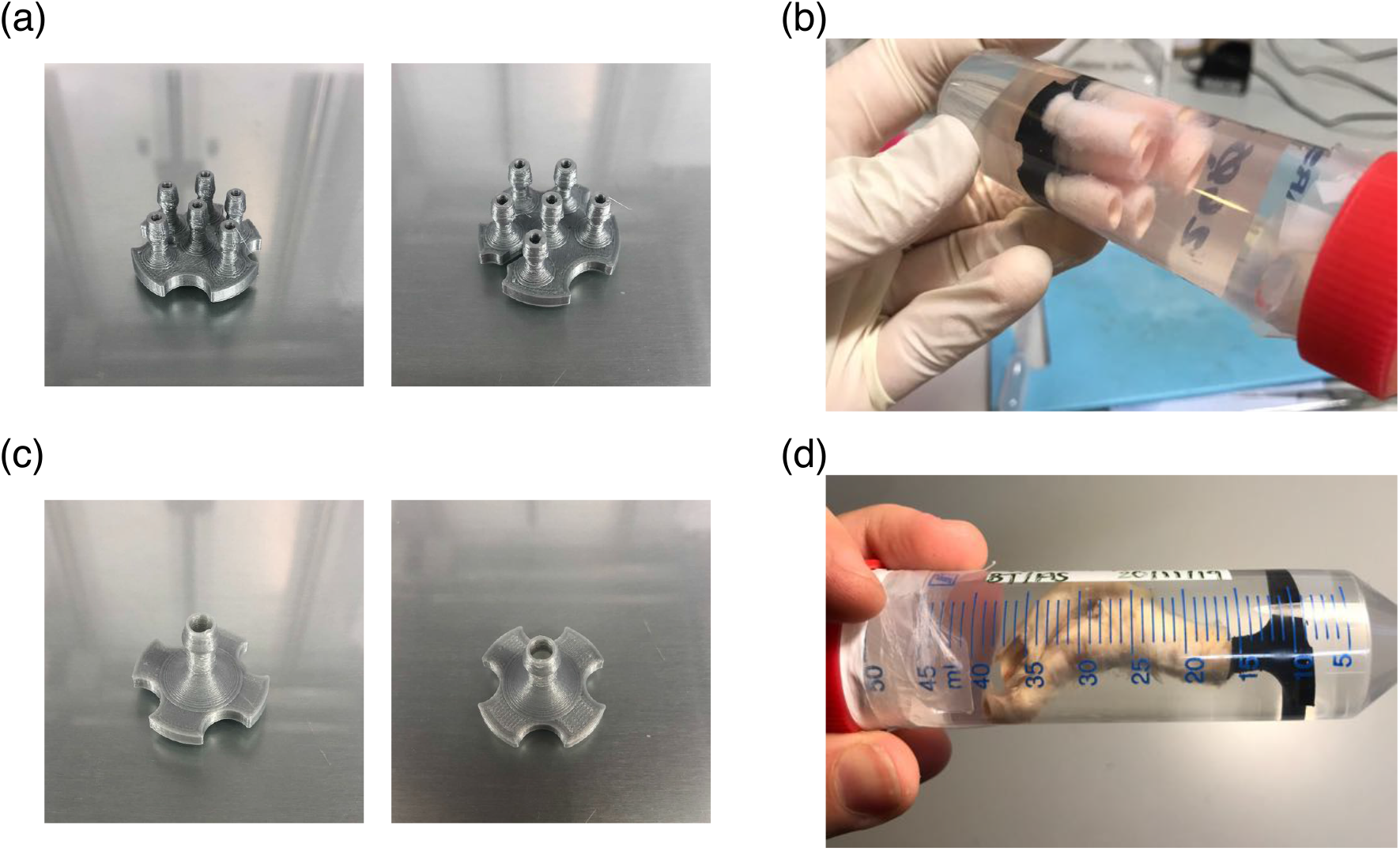
To facilitate imaging of *ex vivo* carotid arteries custom-made 3D printed holders were used for porcine (a) and human (c) carotid arteries. Final sample set-up prior to MR imaging shows porcine (b) and human (d) carotid arteries positioned in 50 ml Falcon tubes and suspended in PBS using the custom holders. Note that 3D printed holders are composed of a bioplastic (polylactic acid) and appearance varies due to the use of poly lactic acid filaments with different colours – silver coloured filament (a) and (c), black coloured filament (b) and (d).

**Figure S2:**
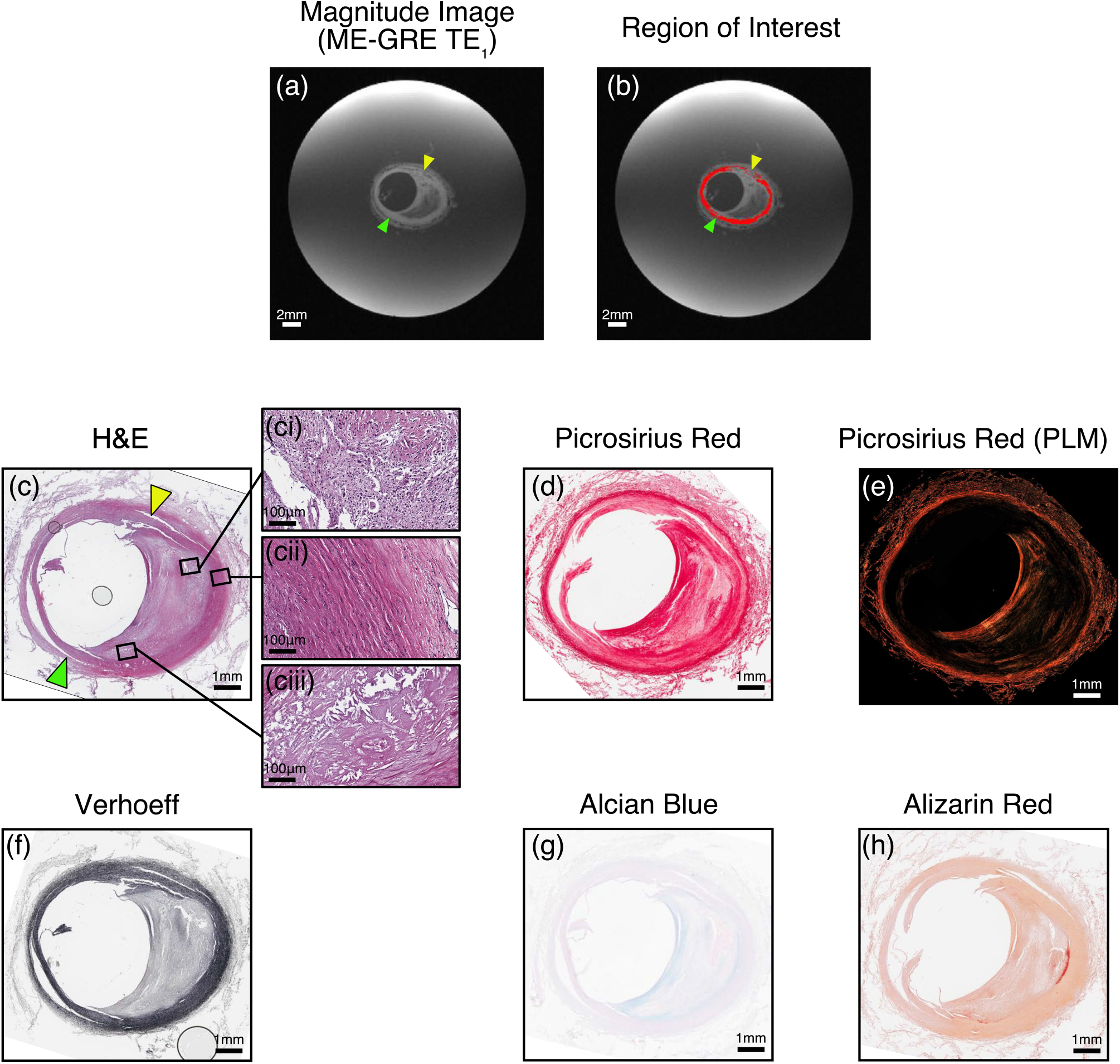
Magnitude image (a) and common carotid ROI definition (b) informed by histology for cadaver sample S01. ROIs were limited to normal appearing arterial tissue in the common carotid. Regions of abnormal smooth muscle cells (c), collagen (d and e), elastin (f), GAG (g) and calcium (h), as identified by histology, were excluded from the final ROI. From H&E histology regions of cell debris indicative of necrotic core (ci), normal appearing tissue (cii) and the cholesterol crystals (ciii) are evident

**Figure S3:**
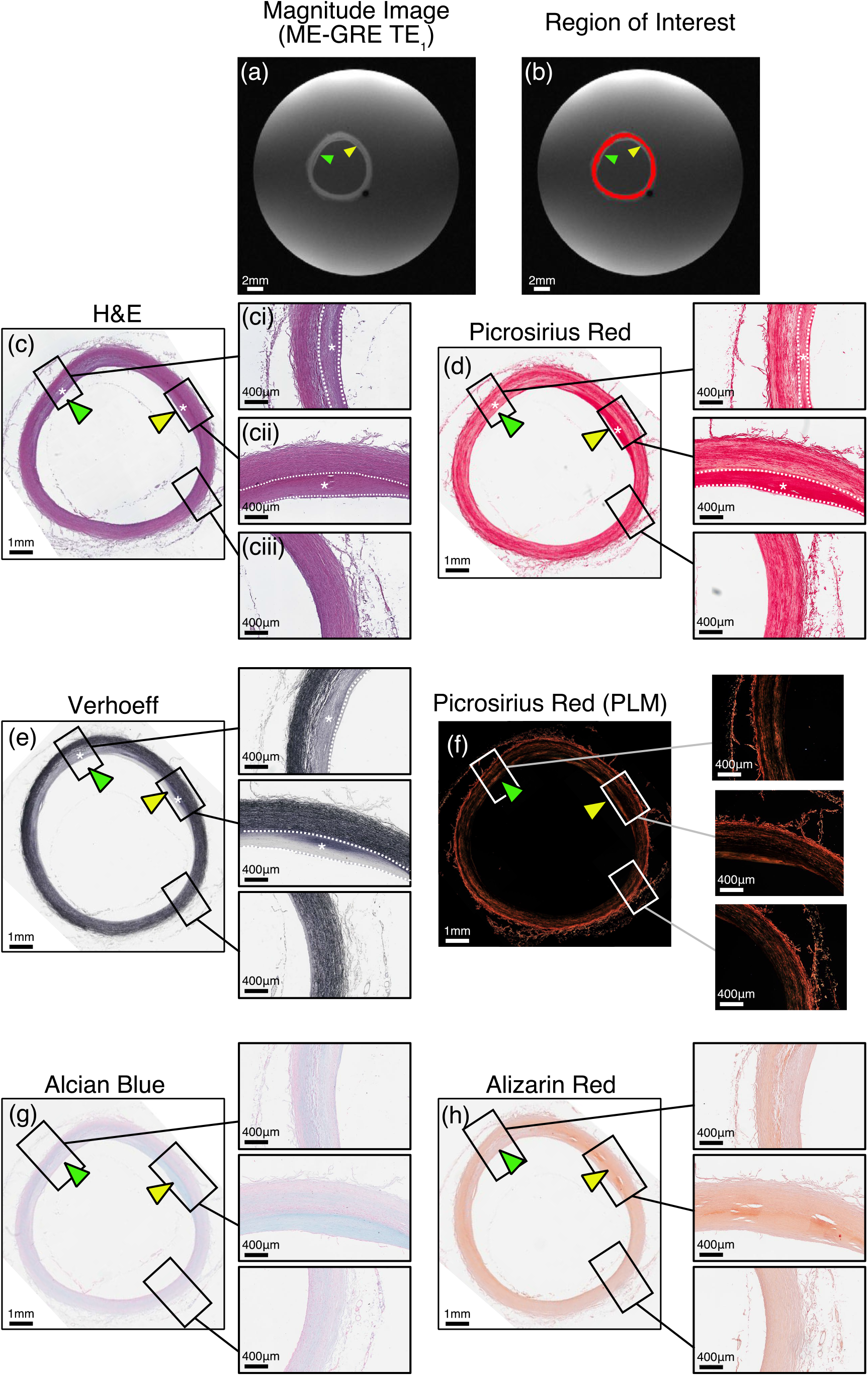
Magnitude image (a) and common carotid ROI definition (b) informed by histology for cadaver sample S02. ROIs were limited to normal appearing arterial tissue in the common carotid. Regions of abnormal smooth muscle cells (c), collagen (d and f), elastin (e), GAG (g) and calcium (h), as identified by histology, were excluded from the final ROI. From H&E histology regions of thickened intima (ci and cii) and normal appearing tissue (ciii) are evident. Regions (ci) and (cii) were excluded from the final ROI

**Figure S4:**
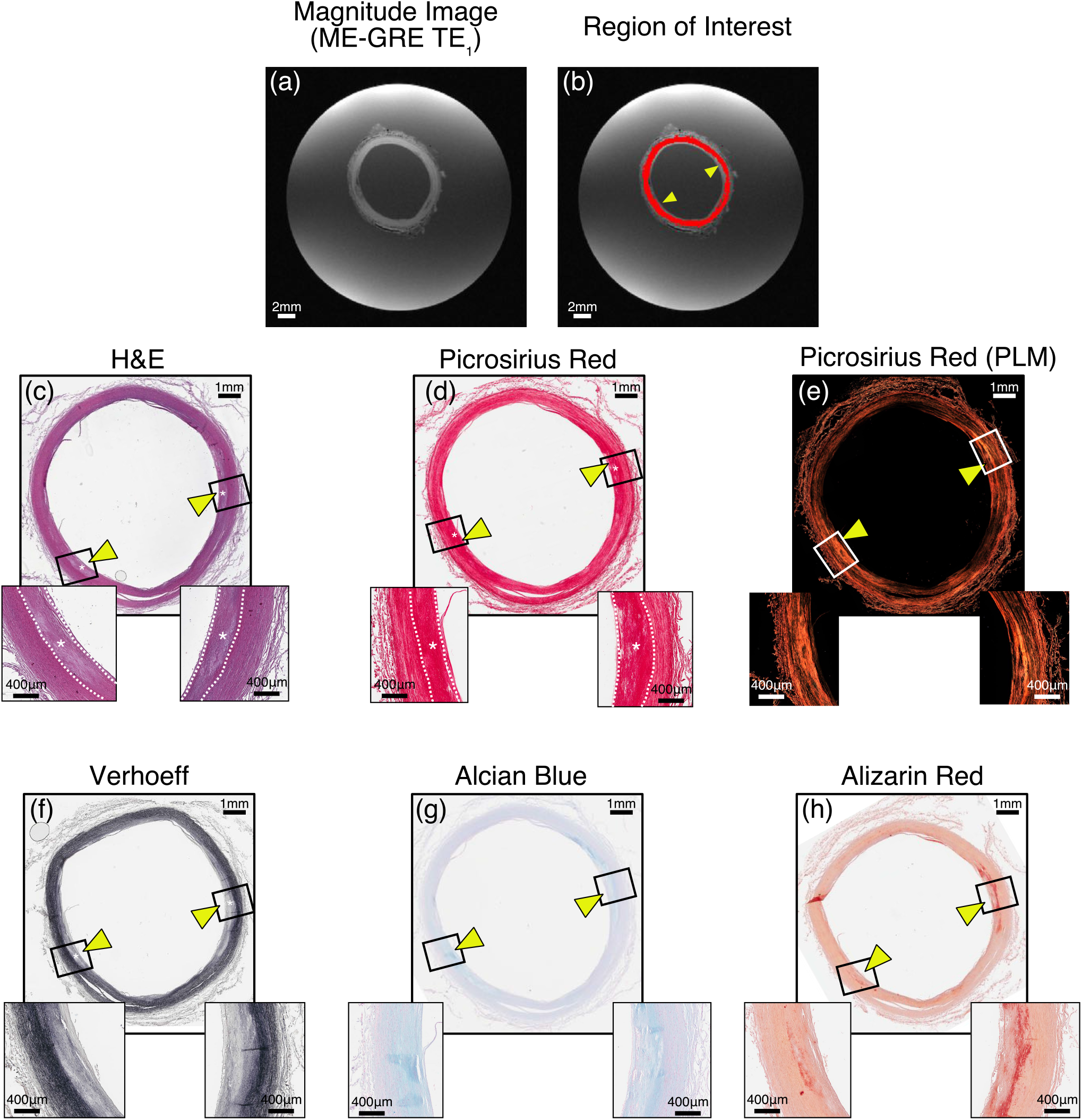
Magnitude image (a) and common carotid ROI definition (b) informed by histology for cadaver sample S03. ROIs were limited to normal appearing arterial tissue in the common carotid. Regions of abnormal smooth muscle cells (c), collagen (d and e), elastin (f), GAG (g) and calcium (h), as identified by histology, were excluded from the final ROI. From histology, bilateral regions of thickened intima are evident and excluded from the final ROI (yellow arrows)

**Figure S5:**
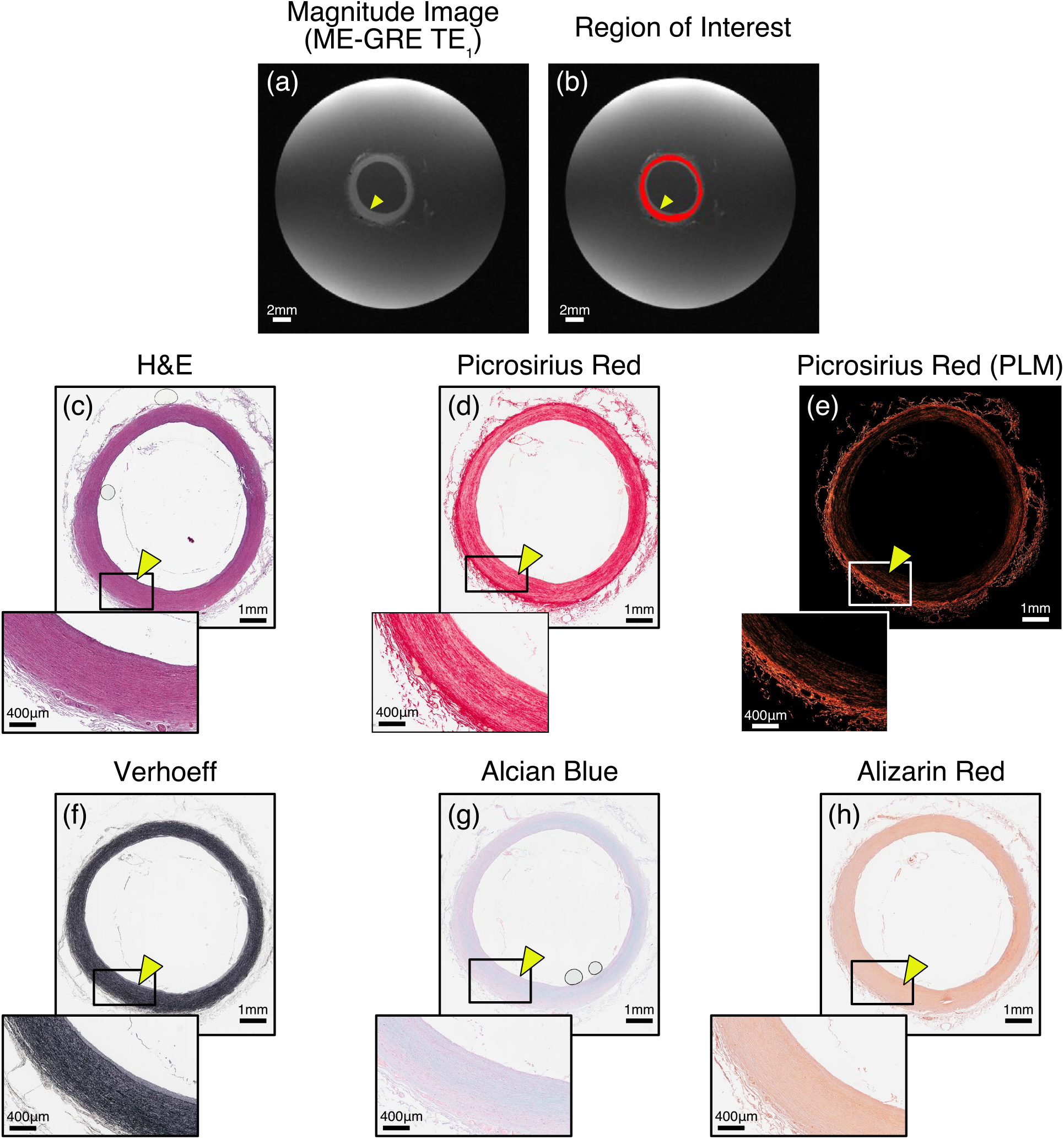
Magnitude image (a) and common carotid ROI definition (b) informed by histology for cadaver sample S04. ROIs were limited to normal appearing arterial tissue in the common carotid via histology visualising smooth muscle cells (c), collagen (d and e), elastin (f), GAG (g) and calcium (h)

**Figure S6:**
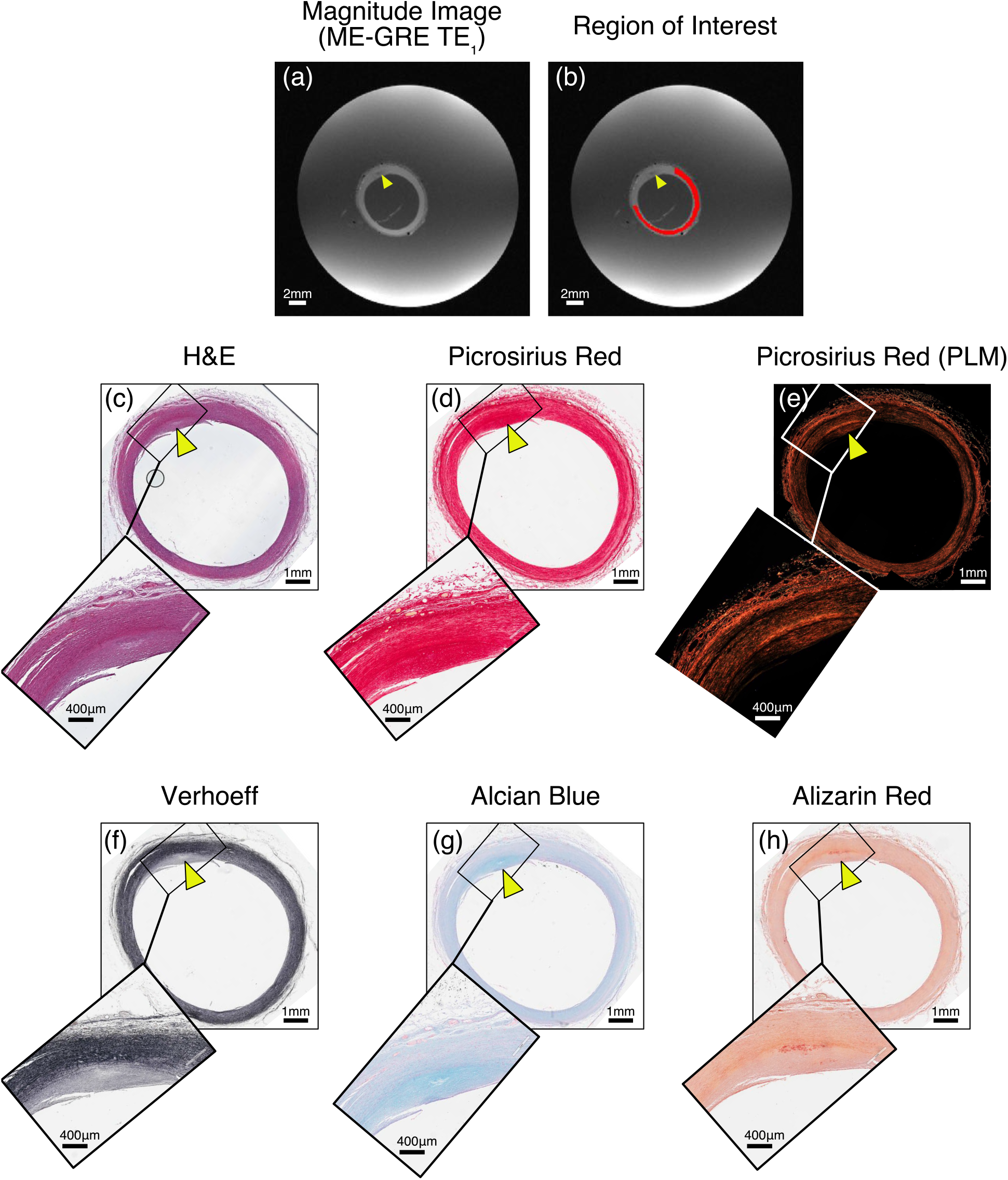
Magnitude image (a) and common carotid ROI definition (b) informed by histology for cadaver sample S05. ROIs were limited to normal appearing arterial tissue in the common carotid. Regions of abnormal smooth muscle cells (c), collagen (d and e), elastin (f), GAG (g) and calcium (h), as identified by histology, were excluded from the final ROI. From histology, a region of thickened intima is evident and excluded from the final ROI (yellow arrow)

**Figure S7:**
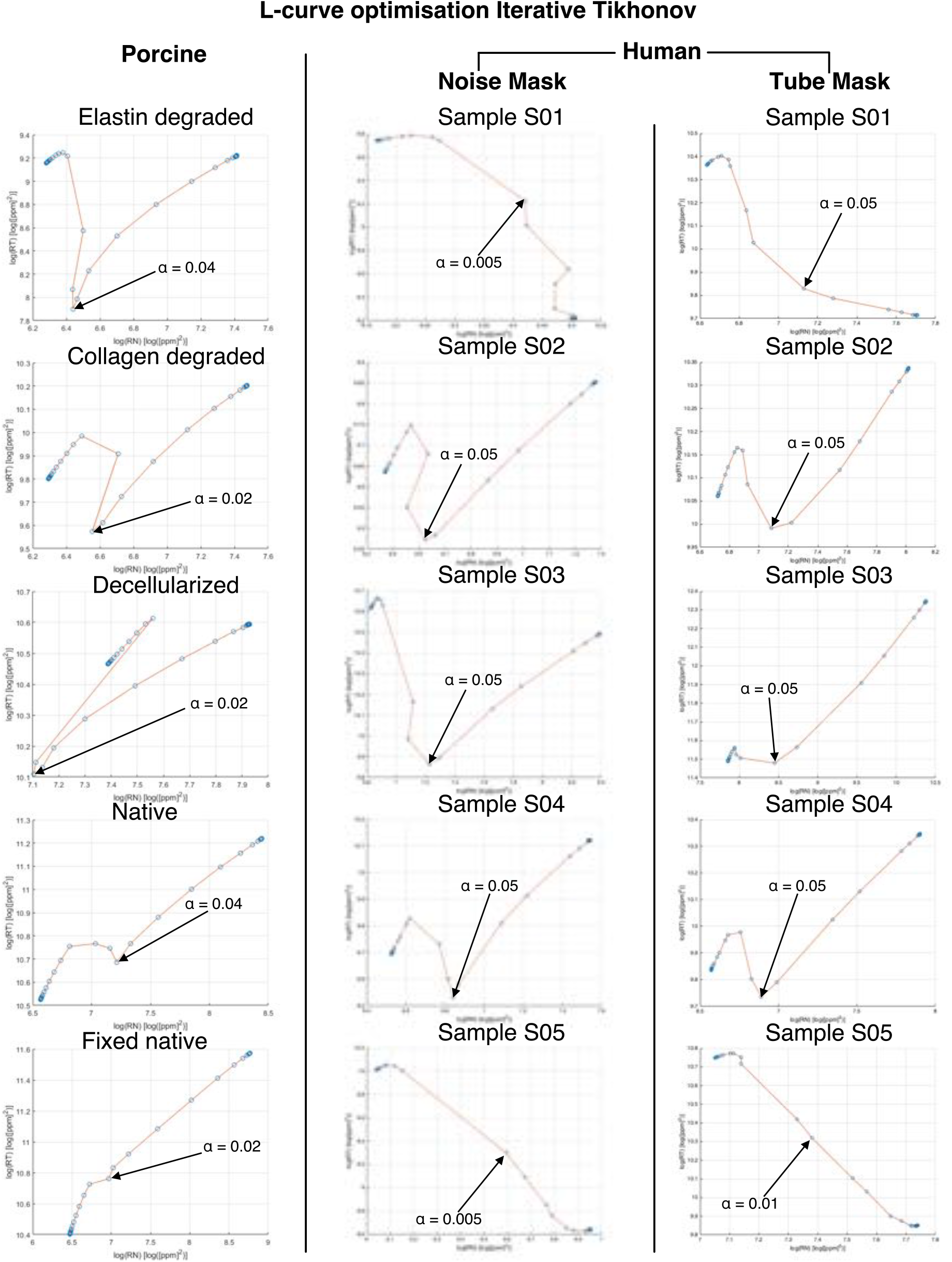
L-curves of all samples presented in this study used for choosing iterative Tikhonov regularisation parameter, α. The optimal iterative Tikhonov regularisation parameter was determined to be α=0.02 for this study

**Figure S8:**
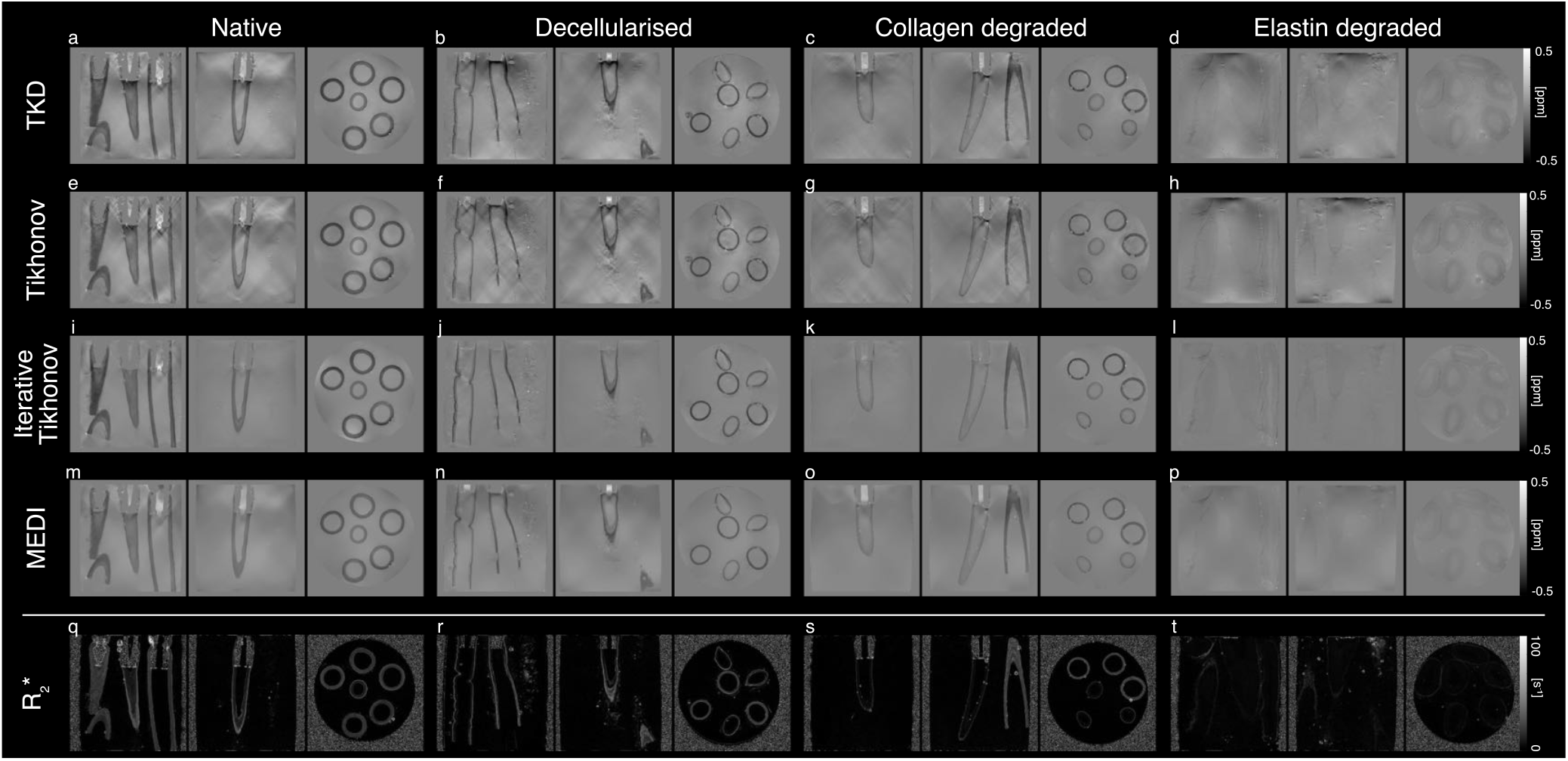
Susceptibility maps produced for the different porcine carotid tissue models (native, decellularised, collagen degraded and elastin degraded) using different susceptibility calculation approaches (TKD, direct Tikhonov, iterative Tikhonov and MEDI). Maps of R_2_* presented on bottom row

**Figure S9:**
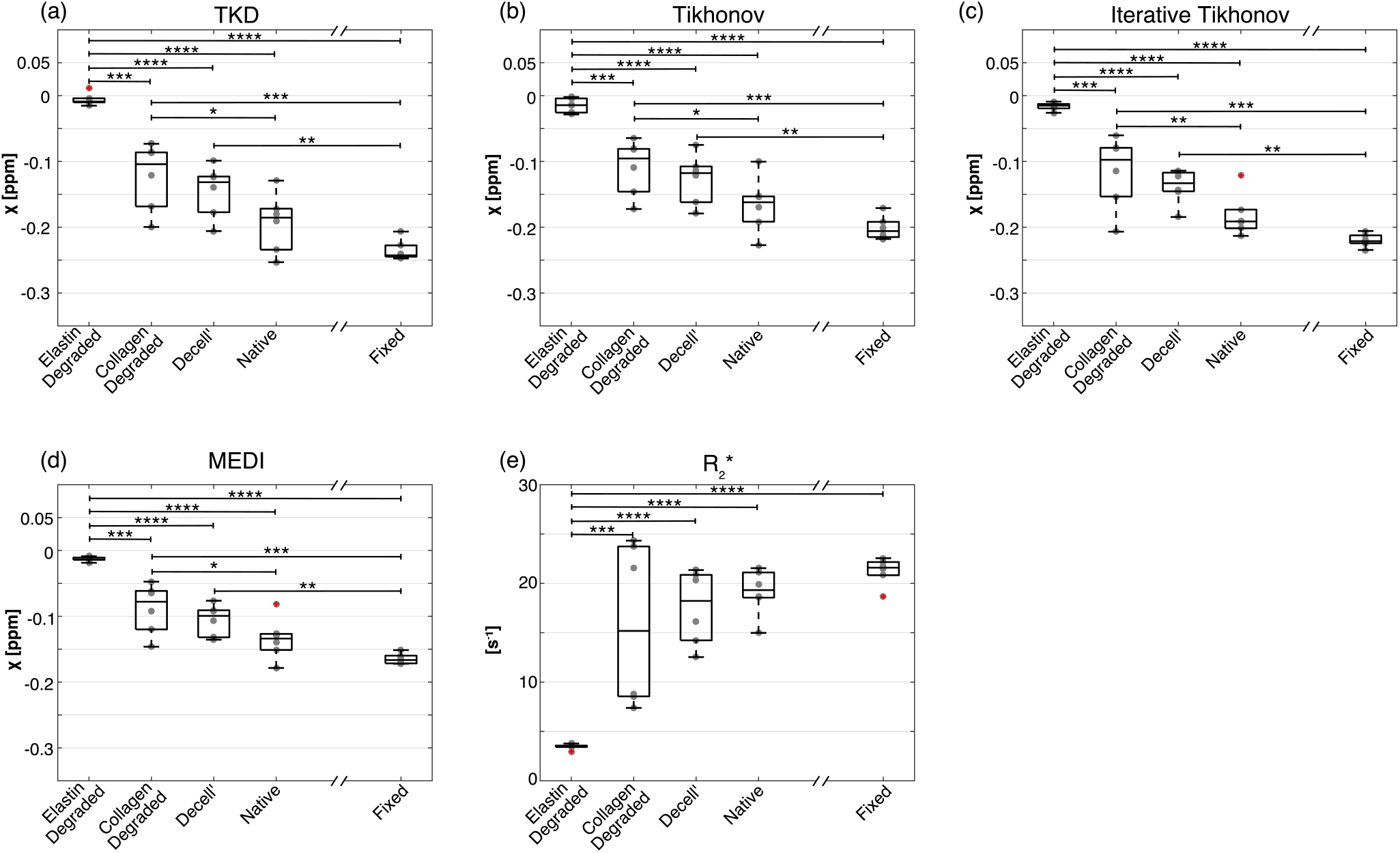
Boxplots comparing susceptibility measurements extracted from vessels (n = 6) of each tissue model for different susceptibility calculation approaches (TKD (a), direct Tikhonov (b), iterative Tikhonov (c) and MEDI (d)). Susceptibility values for tissue models compared using ANOVA and post-hoc test (* p < 0.05, ** p < 0.01, *** p <0.001, **** p < 0.0001)

**Figure S10:**
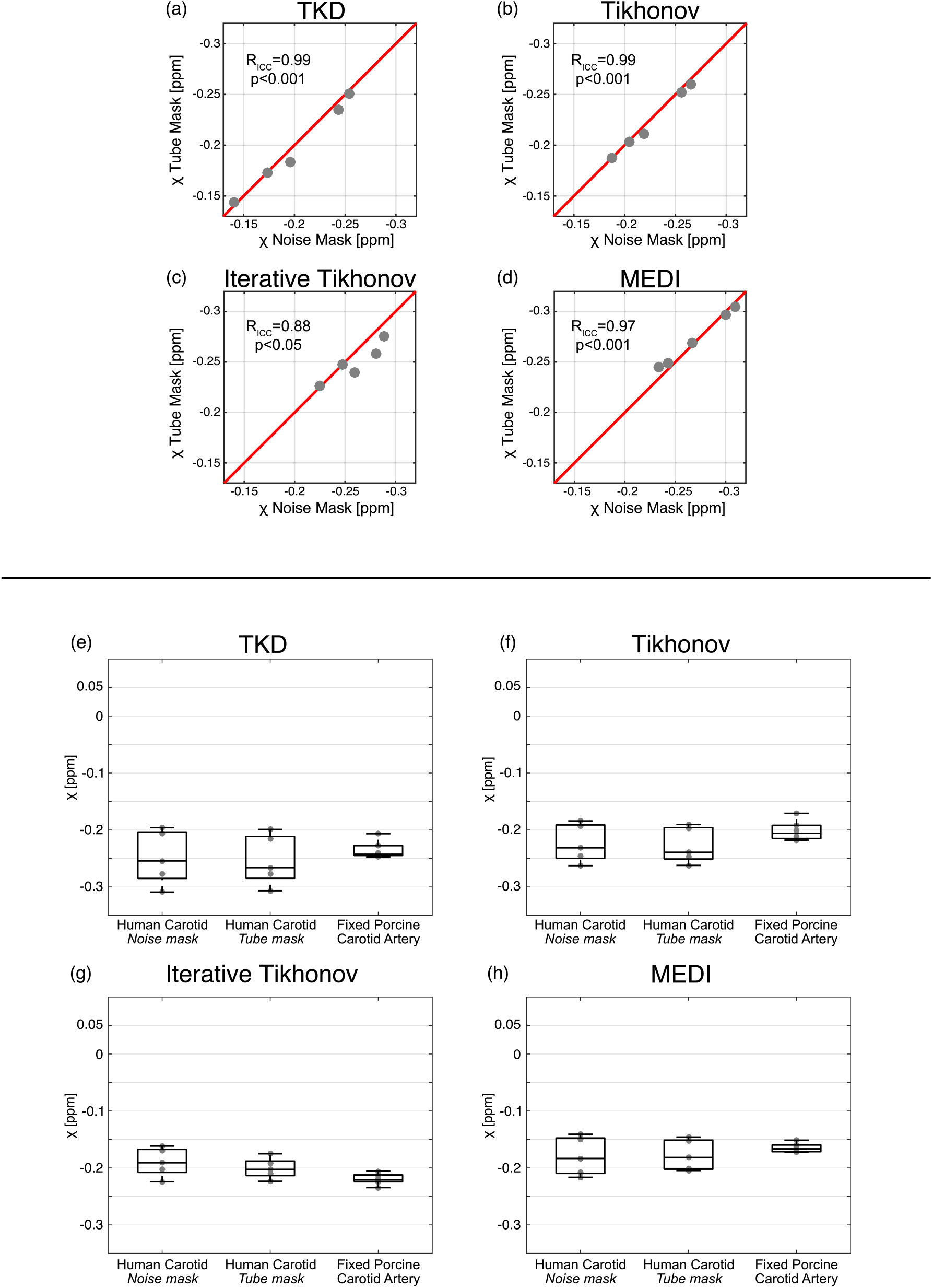
(**a-d**) Comparison between masking pipelines of susceptibility measures in cadaver common carotid. For all dipole inversion approaches, agreement between masking pipelines was excellent using the following criteria - Poor < 0.4, Fair - 0.41– 0.59, Good - 0.60– 0.74, Excellent > 0.75 [Ref 50 main text]. (**e-h**) Boxplots comparing susceptibility measurements in cadaver common carotid artery with fixed porcine carotid artery. Susceptibility values were compared using ANOVA and no significant difference was seen between groups (TKD p=0.7197, Tikhonov p=0.27322, Iterative Tikhonov p=0.050082 and MEDI p=0.559292)

